# Connecting crowns: Analyzing morphological covariation in the modern human postcanine dentition

**DOI:** 10.1101/2024.09.05.611460

**Authors:** Petra G. Šimková, Viktoria A. Krenn, Cinzia Fornai, Lisa Wurm, Vanda Halász, Dominika Lidinsky, Gerhard W. Weber

## Abstract

Morphological covariation within the modern human postcanine dentition remains an open field of study. Analysis of covariation patterns of the three-dimensional (3D) shape between different tooth types has been seldom conducted, but it is relevant for the advancement of human biology and evolution, as well as dental anthropology, phylogeny, and medicine. Here, we analysed 3D shape covariation of the postcanine dentition (excluding third molars), both within and between dental arches using geometric morphometrics (GM). Based on high-resolution (µCT) scans of 526 teeth from 136 individuals we found high pairwise correlation in tooth pairs within the dental arches (lower P3 and P4, r_1_ = 0.89; upper P3 and P4, r_1_ = 0.81; upper M1 and M2, r_1_ = 0.86). The correlation values between antagonists varied notably from the highest value detected between upper and lower M1s (r = 0.9), to the lowest between upper P4s and lower M1s (r = 0.58). Of all analysed tooth types, only the upper M1s showed moderate to high correlation in every pair analysis. Noticeably, unusually high covariation was detected between some of the tooth type pairs that do not articulate in a normal dentition (e.g., lower P3 and upper M2, r_1_ = 0.88). Furthermore, a relatively high covariation was found in the pairs of lower P4s and M1s (r_1_ = 0.79), and upper P4s and M1s (r_1_ = 0.77), which are the only tooth type pairs of the postcanine dentition belonging to different tooth classes (premolars and molars, respectively) and still serving similar masticatory functions.

This study points to the fact that higher morphological integration seems to characterize teeth within the same dental arch rather than between antagonistic teeth. With this study, we provided an overview of pairwise correlations and strength of covariation between different tooth types. This information might inform future studies aimed at understanding developmental, phylogenetic, and functional aspects of the human postcanine dentition, including possible phenotype-genotype associations. However, with this study being the first one performed on a 3D sample of this size, we also report on obstacles and peculiarities that have been determined.

## Introduction

Human dental morphology has been a topic of interest in the fields of dental medicine, biological anthropology, and human evolution since the last century, with a particular focus on exploring dental geometry of various tooth types in 3D, an approach emerging in the past two decades (Bailey et al., 2011; Benazzi et al., 2012, 2014; Fornai et al., 2014; Hershkovitz et al., 2018; Krenn et al., 2019; Macchiarelli et al., 2013; Monson et al., 2020; Morita et al., 2014, 2016; Ortiz et al., 2017; Pan et al., 2016; Šimková et al., 2021, 2024; Weber et al., 2016; Zanolli et al., 2018). Since the 3D crown shapes of the human postcanine dentition have never been analyzed in relation to each other, the phenotypic covariation of the actual geometry of human postcanine teeth is still unknown, yet crucial to be connected to the genotype in a further step. Research on developmental genetics has linked dental genotype to phenotype (Hlusko, 2016; Thesleff, 2014), but while the odontogenetic morphological stages are well-documented (Thesleff & Tummers, 2008), the genetic pathways regulating odontogenesis remain under investigation. Quantitative genetics distinguishes populational phenotypic variation effects into genetic, non-genetic, and variation due to covariate effects. Similarly, phenotypic correlations can be broken down into genetic and non-genetic mechanisms, enabling the exploration of genetic covariation patterns underlying phenotypic traits (Hlusko, 2016). The comprehensive study of dental phenotypic variation in 3D instead of traditional linear distances can represent a major step forward in identifying morphological patterns and their underlying genetic mechanisms.

The development of anatomical structures is a coordinated process driven by growth factors, induction processes, signaling molecules and cascades, in which adjacent elements interact in order to create tightly integrated forms. This process, referred to as developmental integration, synchronizes the formation of various tissues, generating cohesive patterns of phenotypic variation throughout growth. Apart from developmental integration, pleiotropy (genes affecting multiple traits; Cheverud, 1996; Hlusko & Mahaney, 2009; Hodgkin, 1998; Wagner & Zhang, 2011) as well as linkage disequilibrium (deterministic association of alleles on multiple loci, with an effect on various traits; Mitteroecker et al., 2012) affect the induction of phenotypic covariation in populations. The need for a tight link between anatomical elements that co-participate in a particular function (functional integration), leads to correlational selection that can result in linkage disequilibrium, and thus in covariation between structures that are not developmentally linked. However, even with robust selection pressures, the effect on phenotypic covariation is lower in comparison to the influence of developmental integration and pleiotropic effects (Mitteroecker et al., 2012).

More than 2400 genes appear to be involved in odontogenesis (Landin et al., 2012; Sehic et al., 2017), with the majority of related proteins belonging to four signaling pathway families; BMP, FGF, SHH, and WNT (Bei, 2009; Hosoya et al., 2020; Jussila & Thesleff, 2012; Mikkola, 2007; Yuan & Chai, 2019), as well as to the proteoglycans (Chen et al., 2023). Various evolutionary developmental models have been proposed to explain odontogenesis (Butler, 1939; Jernvall, 2000). Kavanagh et al. (2007) significantly contributed to the inhibitory cascade model (which describes how the development of one tooth might influence the adjacent teeth), demonstrating that activator-inhibitor dynamics determine size differences between molars and significantly contribute to metameric variation within a tooth row (Morita et al., 2016). Resulting from both neutral and adaptive processes, various parts of the primate dentition are heritable and vary significantly inter-as well as intra-specifically (Hanihara & Ishida, 2005; Hlusko et al., 2011; Monson et al., 2019, 2020). A high genetic contribution to dental morphological variation in humans has also been underlined by twin studies (Hughes et al., 2014; Townsend et al., 2012). In an experimental study on *Cryptotis parva’s* molars, narrow-sense heritability (proportion of total phenotypic variance that is attributed to additive genetic variance alone) of overall crown shape has been determined at h^2^=0.34 (in a range between 0 and 1), thus playing an important role in determining the shape of the tooth crown (Polly & Mock, 2018). However, as mentioned above, covariation between morphological traits is not only influenced by shared developmental and genetic factors but also by variations in the growth processes and allele frequencies that underlie it (Mitteroecker et al., 2012). The concept of morphological integration, where phenotypic traits are correlated if they share a developmental pathway or function, and modularity (Wagner & Altenberg, 1996) that explains the mechanisms behind morphological integration, are the supporting concepts of this approach. Modules are further classified as developmental, functional, genetic or evolutionary (Klingenberg, 2008), and can influence each other, causing phenotypic covariation (Cheverud, 1996). Assessment of the covariation patterns within the human dentition is crucial for understanding odontogenesis and modularity (Hlusko et al., 2011; Stock, 2001). Genetic studies showed that odontogenesis involves hierarchical genetic patterning mechanisms across and between dental arches, as well as within and between tooth types (Jernvall & Thesleff, 2000). One of the possible investigation approaches uses genetic and genomic analyses to identify loci linked to dental variation (Geller et al., 2011; Pillas et al., 2010), though these methods do not clarify the mechanisms behind morphological variation.

Functional occlusion is key for efficient food breakdown and it relies heavily on the complementary 3D topography of the occluding dental surfaces (antagonists; Nelson & Ash, 2010). Therefore, we can expect the morphological shape correlation between antagonistic teeth to be high, while odontogenetic mechanisms suggest higher shape covariation within dental arches. This study explored the shape covariation between modern human postcanine dentinal crowns (third molars excluded) within and between upper and lower dental arches working by means of geometric morphometrics (GM) and 3D imaging. To our knowledge, no research has yet used comprehensive 3D dental shape data to explore the covariation between both third and fourth premolars, as well as the first and second molars of both jaws. In this project we selected high-resolution (µCT) scans of 526 teeth from 136 individuals for analyses based on the condition of their dentition, ensuring the representation of as many dental types per individual as possible. The focus of our approach was on the enamel-dentine junction (EDJ) that is the primary developmental structure of the crown and is consequently the precursor of the enamel cap’s morphology, which naturally results in a strong correlation between EDJ and the outer enamel surface (OES; Bailey et al., 2011; Fornai et al., 2015; Guy et al., 2015; Monson et al., 2020). The EDJ thus represents an appropriate alternative to the OES (Gómez-Robles et al., 2013; Skinner et al., 2008, 2009; Weber et al., 2016) and since the overlying enamel protects the EDJ from early deterioration processes such as abrasion, erosion or chipping, its examination offers advantages. This approach will enable us to analyze every dental type within and between dental arches in most of the individuals and explore potential differences between human populations.

## Materials and Methods

Altogether, this study investigated 526 teeth from the upper and lower postcanine dentitions of 136 individuals from seven geographically different human groups (Egyptians, n = 26, Europeans, n = 123, Near Easterners, n = 69, Oceanians, n = 59, Sub-Saharan Africans, n = 109, South Americans, n = 83, and Southeast Asians, n = 57). In particular, we considered teeth from eight different tooth types, namely 70 lower third premolars (P3s), 74 lower fourth premolars (P4s), 76 upper P3s, 74 upper P4s, 49 lower first molars (M1s), 48 lower second molars (M2s), 59 upper M1s, and 76 upper M2s. We use the term “tooth class” to refer to premolars or molars in general. We relied on the study of osteological specimens from museum collections since they present natural dental wear, rarely show any caries, and mostly no dental intervention such as fillings. Importantly, our study required µCT images which cannot be obtained from living subjects. However, assembling a sample with fully preserved postcanine dentition from osteological collections was challenging because of missing, broken or severely worn teeth, resulting in differing subsample sizes for the various tooth types. For instance, M1s, as the first permanent teeth to erupt, are more prone to wear than other tooth types and are the least represented in our sample.

### Scanning and Segmentation

All specimens were scanned at the Vienna μCT Lab in Austria using an industrial Viscom X8060 NDT scanner with parameters of 110–140 kV, 280–410 mA, 1400–2000 ms, and a 0.75 mm copper filter, achieving a voxel size between 20 and 50 μm. Virtual segmentation of the μCT data was conducted using Amira software (www.thermofisher.com) to separate the dental tissues from each other and from the surrounding area, and subsequently generate triangulated surface models of the enamel and dentine. The models were derived from both the right and left side, but the datasets representing right teeth were flipped prior to surface extraction to avoid later reflection of the landmark coordinates. Teeth are not expected to exhibit directional asymmetry (Frederick & Gallup, 2007), thus we considered this procedure uninfluential on the outcomes of our study. In cases of slightly worn dentinal horn tips (maximal stage 3; Molnar, 1971), virtual reconstruction was performed in Amira using the ‘brush’ tool by extending the existing contours of the dentinal horn tips.

### Reorientation and Data Collection

The segmented 3D surface models were reoriented in Geomagic Design X 64 (www.3dsystems.com) following established protocols according to the dental type. The cervical plane, calculated as the best-fit plane of the cervical margin, was used to reorient the dataset parallel to the x-y plane of the virtual workspace. The crowns were then rotated to align the upper premolars’ and molars’ mesial ridge parallel with the y-axis, the lower premolars’ buccal aspect of the marginal ridge with the x-axis and the lower molars’ lingual aspect of the cervical outline with the x-axis. The cervical outlines of the reoriented surfaces were collected and projected onto the cervical plane. These outlines were imported into Rhinoceros 6 (www.rhino3d.com) and divided into 24 segments using equiangular radial vectors originating from the area centroid of the outline. Subsequently, 24 pseudo-landmarks were placed at the points of intersections between the outline and the radial vectors, representing the cervical outline of every tooth type.

### Landmark Collection on the EDJ

The reoriented 3D surfaces of the dentinal crowns were imported into the EVAN-Toolbox software (www.evan-society.org) for landmark collection, using configurations suitable to the different dental types (Fig. 1). For all premolars, two fixed landmarks (LMs) were placed on the paracone and protocone horn tips. On both lower premolars (Krenn et al., 2019), as well as, on the upper P3s, two additional LMs were placed at the deepest mesial and distal points of the central groove. In upper P4s, where the central groove is less defined, the landmarks were instead placed on the deepest points of the distal and mesial ridges (Hershkovitz et al., 2018; Šimková et al., 2024). Additionally, 20 curve semilandmarks (sLMs) were placed along the marginal ridge.

**Fig. 1.**
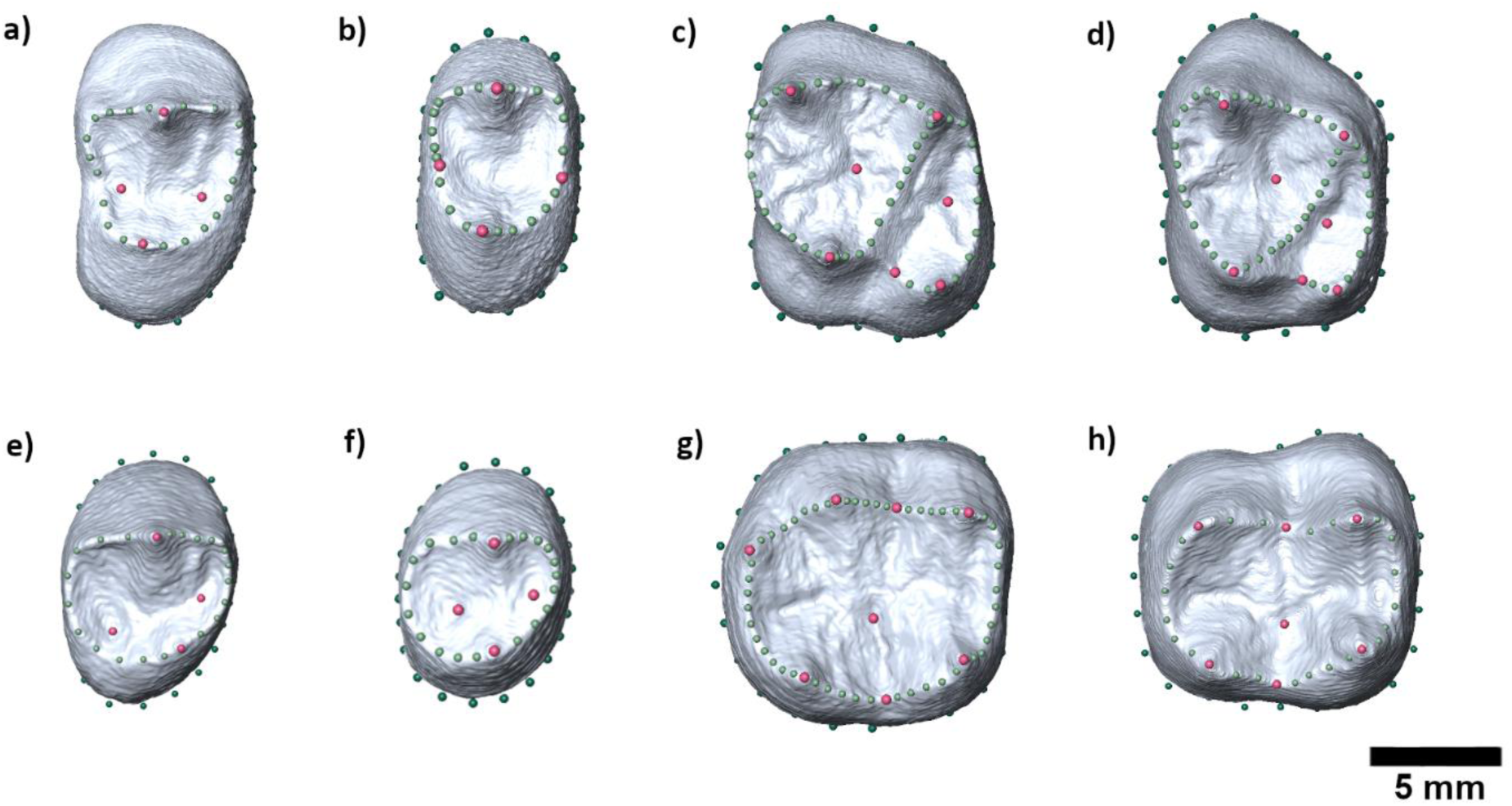
Landmark configurations of the dentinal crown dataset of upper third **(a)** and fourth **(b)** premolars, upper first **(c)** and second **(d)** molars, lower third **(e)** and fourth **(f)** premolars, and lower first **(g)** and second **(h)** molars. Fixed landmarks = pink; semilandmarks = light green; pseudo-landmarks = dark green.

Upper molars were represented by 4 LMs on each of the main cusp’s horn tips, and 3 more LMs placed at the deepest points of the central fossa, distal fossa and on the lowest point between the hypocone and the protocone along the marginal ridge of the EDJ (Hershkovitz et al., 2018). Forty-three sLMs were used to capture the curve of the EDJ in both upper molars.

On lower molars, LMs were placed on the tips of the main horns, including the hypoconulid in the LM1s, as well as on the lowest point of the lingual and buccal marginal ridge, and the deepest point of the central fossa (Weber et al., 2016). The marginal occlusal ridge was represented by 47 sLMs in the LM1s and 22 sLMs in the LM2s.

### Geometric Morphometric Analyses

The shape of the dentinal crowns was analyzed in the EVAN-Toolbox 1.75 software (www.evan-society.org) using the landmark configuration representing both the cervical outline and the EDJ. For each of the subsamples, General Procrustes Analysis (GPA; Gower, 1975) was conducted to normalize the landmark configurations. The obtained Procrustes shape coordinates were then analyzed using a 2-block Partial Least Squares (2B-PLS) analysis to explore shape covariation and pairwise correlation between different dental types of the postcanine dentition within and between the jaws. In this study, PLS Mode A (a symmetric relation between blocks; Rosipal & Krämer, 2006) was employed. Consequently, we did not report on the RV coefficient, as its accuracy can in this case be compromised by factors related to sample size and the number of variables considered (Adams, 2016). Instead, we present the total squared covariance and the pairwise correlations, which provide a standardized measure easy to compare.

## Results

The percentages of total covariance according to the Singular Warp Score 1 and pairwise correlations are listed in Table 1 and Table 2. We found a high pairwise correlation (Table 2) in all pairs of the same tooth class within the dental arch (lower P3s and P4s, r_1_ = 0.89; upper P3s and P4s, r_1_ = 0.81; upper M1s and M2s, r_1_ = 0.83), except for the lower M1s and M2s that expressed a moderate pairwise correlation of 0.73. The correlation values between antagonists varied notably from very high values calculated for the pair of upper/lower M1s (r_1_ = 0.90) and upper M1s/lower M2s (r_1_ = 0.82), through moderate values in upper/lower P3s (r_1_ = 0.71) and lower P4/upper P3 (r_1_ = 0.70), to low in upper/lower P4s (r_1_ = 0.65), upper/lower M2s (r_1_ = 0.64) and upper P4/lower M1s (r_1_ = 0.58). Moderate to low correlation values were detected between non-neighboring tooth types located in the same jaw, as well as tooth types that do not touch in full occlusion at all, with most of the relatively high values detected in those tooth pairs that included a lower premolar. An unexpectedly high pairwise correlation was found in the pair of lower P3 and upper M2 (r_1_ = 0.88). A relatively high pairwise correlation was also found in the pairs of lower P4s and M1s (r_1_ = 0.79), and upper P4s and M1s (r_1_ = 0.77), which are the only pair of adjacent postcanine teeth belonging to different dental classes, yet serving similar functional purposes during mastication (Kraus et al., 1969). Additionally, the correlation between the upper P4 and M2 (r_1_ = 0.78) was as strong as with upper M1s. We detected a coherent pattern in the overall shape change of the crowns, consistent in all tooth types, namely the change between tall and narrow crowned teeth versus short and broad crowned (shape-types described by Pfneiszl (2024)). A thorough description of the actual 3D morphological changes of the covarying occluding or neighboring tooth type pairs can be found in the caption of Figure 3 and in the Supplementary Information (Supplementary Fig. S1-S15).

**Table 1.**
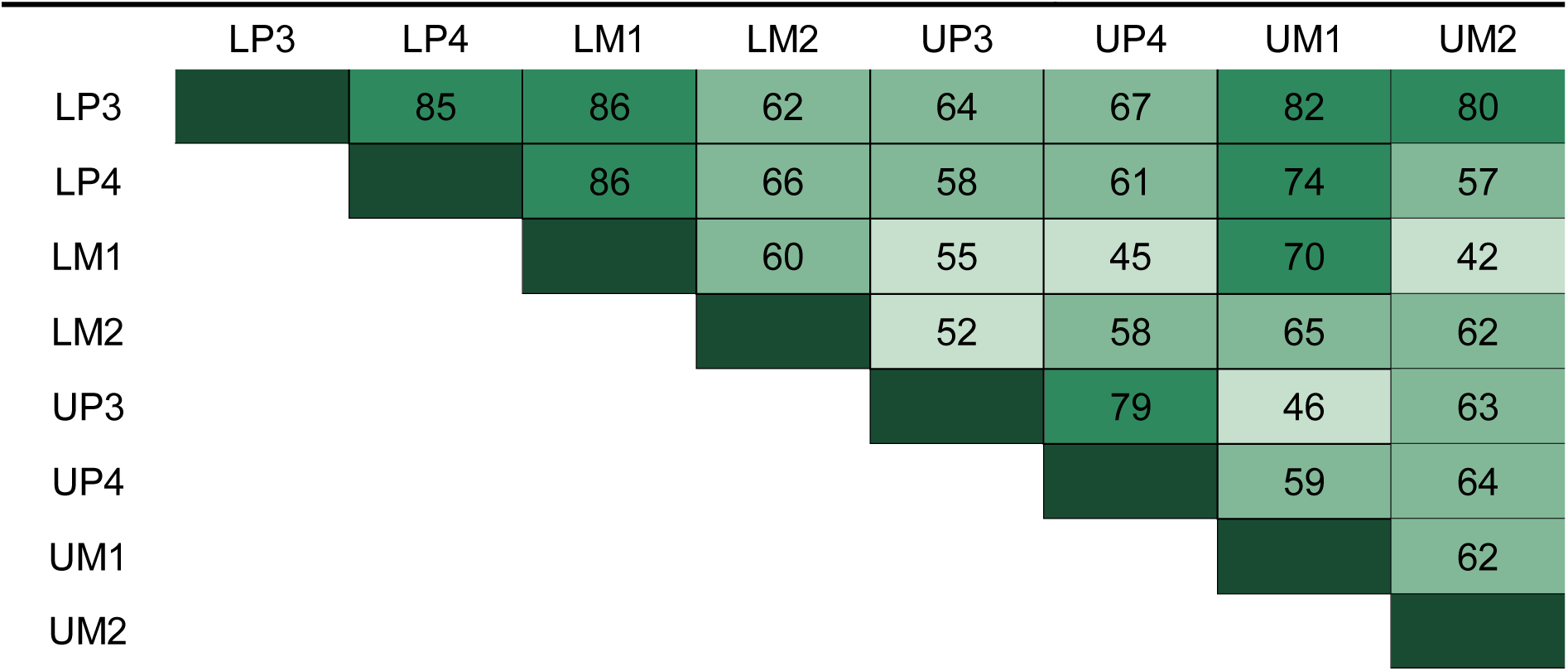
Percentages of total squared covariance for the Singular Warp Score 1, resulting from 2-block Partial Least Squares Analyses. (L = lower; U = upper; P3 = third premolars: P4 = fourth premolars; M1 = first molars; M2 = second molars)

|  | LP3 | LP4 | LM1 | LM2 | UP3 | UP4 | UM1 | UM2 |
| --- | --- | --- | --- | --- | --- | --- | --- | --- |
| LP3 |  | 85 | 86 | 62 | 64 | 67 | 82 | 80 |
| LP4 |  |  | 86 | 66 | 58 | 61 | 74 | 57 |
| LM1 |  |  |  | 60 | 55 | 45 | 70 | 42 |
| LM2 |  |  |  |  | 52 | 58 | 65 | 62 |
| UP3 |  |  |  |  |  | 79 | 46 | 63 |
| UP4 |  |  |  |  |  |  | 59 | 64 |
| UM1 |  |  |  |  |  |  |  | 62 |
| UM2 |  |  |  |  |  |  |  |  |

**Table 2.**
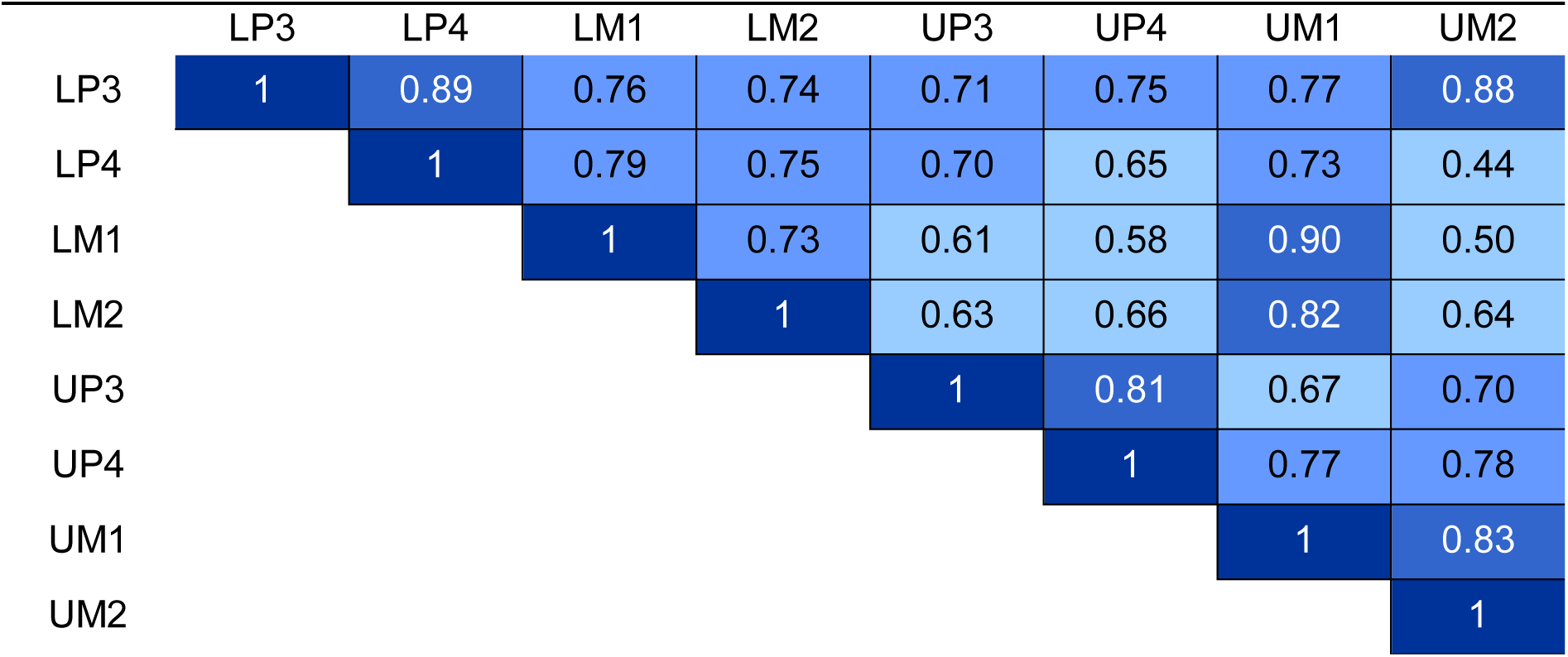
Pairwise correlation values resulting from 2-block Partial Least Squares Analyses. (L = lower; U = upper; P3 = third premolars: P4 = fourth premolars; M1 = first molars; M2 = second molars)

**Fig. 2.**
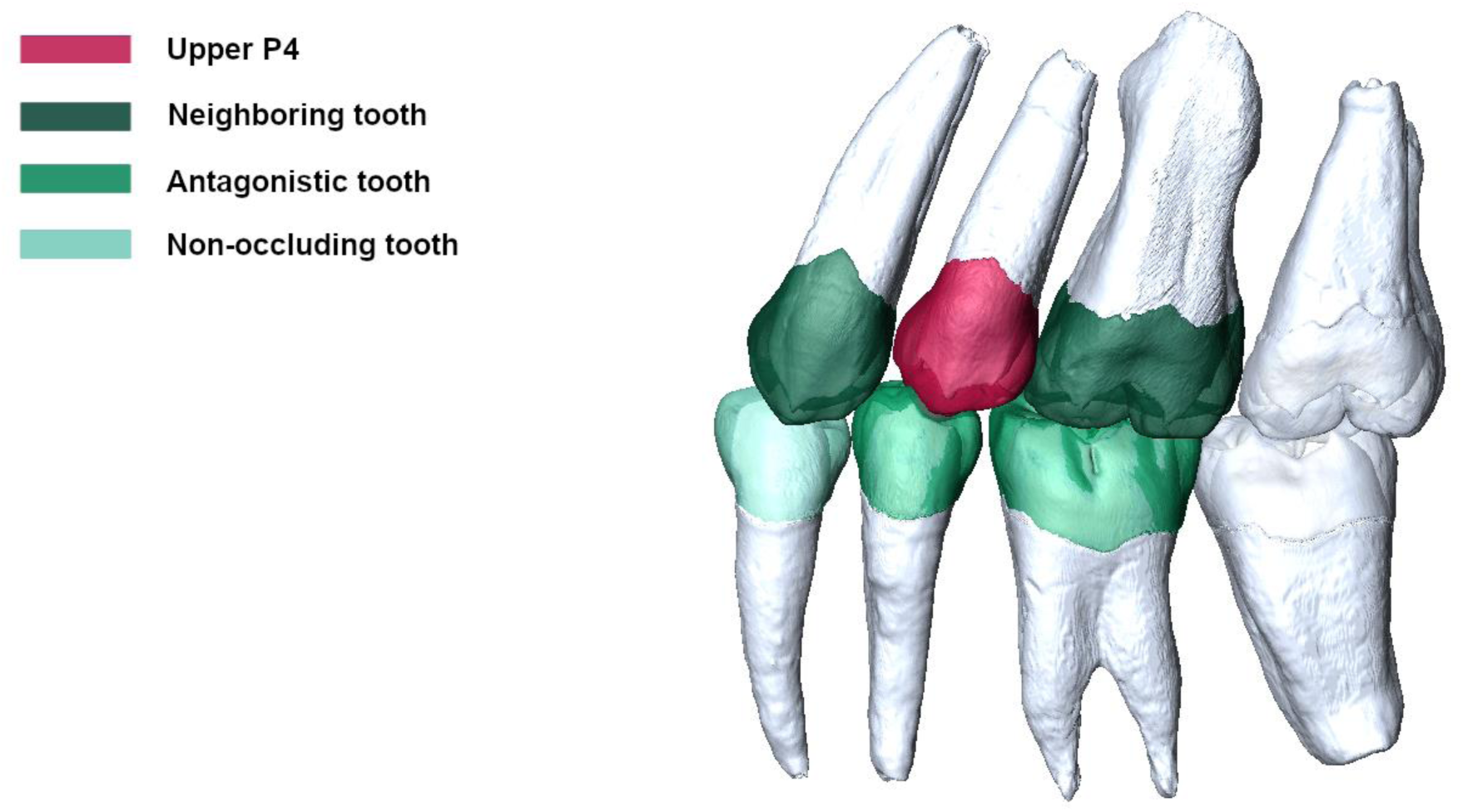
The postcanine dentition (third molars excluded) in occlusion. The color coding is used to describe the specific relationship of the upper fourth molar (chosen as an example) to its neighboring tooth types.

**Fig. 3.**
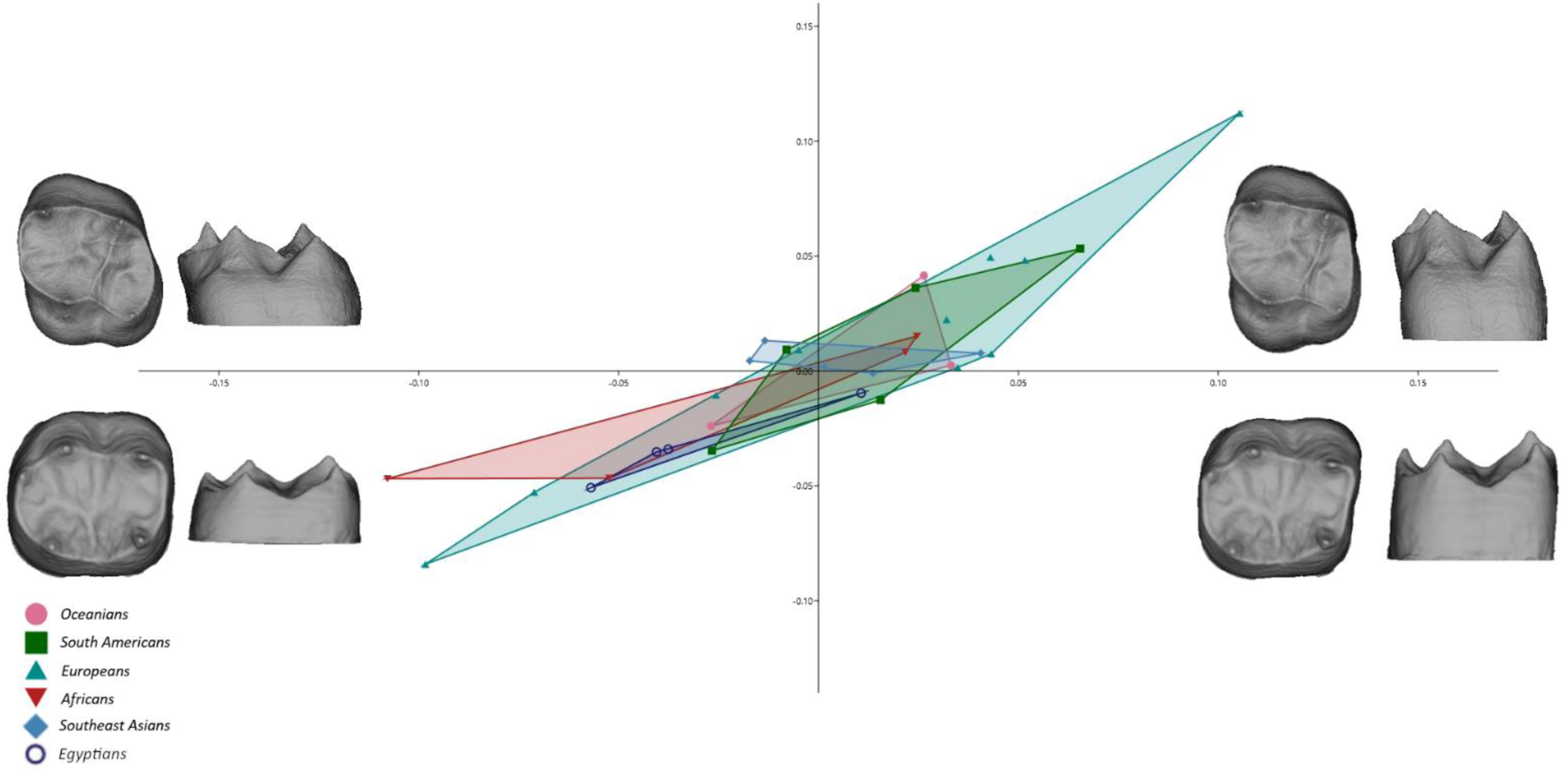
2B-PLS plot for the covariation of upper and lower first molars’ dentinal crowns; the warping shows the real shape variation in occlusal and lingual view at the extremities of the range of distribution. The covariation analysis between antagonistic M1s showed that the main shape change followed the pattern between low and broad crowns versus tall and narrow crowns. In upper M1s, the hypocone was variably expanded relative to the trigon, while lower M1s showed a mesio-distal and bucco-lingual expansion of the cervical outline relative to the occlusal aspect, slightly stronger on the mesial part. The occlusal aspect of the lower M1 remained rather stable, showing more rounded mesial and distal aspects, with the relative expansion of the talon.

Regarding geographical variation in our sample, we found a great overlap of morphological expressions across populations with larger and partially also smaller sub-sample sizes (e.g., Figure 3). However, the analyses that revealed a complete or partial separation between populations, are also those with the smallest sample sizes (see Supplementary Information).

## Discussion

In previous studies, high pairwise correlation has been reported for premolars in the lower jaw (r_1_ = 0.87; Krenn et al., 2019), as well as in the upper jaw (r_1_ = 0.83; Šimková et al., 2024), indicating that the morphology of both upper and lower premolars varies in a concordant manner, with the crown shape-type being either tall and narrow or low and broad. The current study included analyses of extended samples with more individuals from additional human populations which confirmed the results of previous studies. Furthermore, agreeing with the findings of Buchegger et al. (2016), we found that the third highest correlation within this tooth class occurred between lower P3s and upper P4s which do not occlude in normal occlusion (Table 2). Surprisingly, correlations between antagonistic upper and lower P3s and P4s were in contrast lower than expected. This can probably be explained by the significant variability that lower premolars exhibit on the lingual crown side, due to differences in mesial and distal fossae expansion, affecting their lingual cusp positioning (Krenn et al., 2019). Hence, the non-occluding lingual cusp is free to vary, whereas the buccal aspect is more stable, being the most canalized region of this tooth (Siegal & Bergman, 2002). This distinguishes them from their upper antagonists, which are overall more stable and less variable.

Interestingly, P4s covaried more tightly with M1s within the same dental arch, rather than with their antagonists of the same tooth class. Premolars are located between the anterior teeth and the molars and serve distinct functions. In normal occlusion, the buccal cusp of the lower P3 slides across the interproximal embrasure of the upper canine and third premolar, leading Kraus et al. (1969) to consider them a “functional entity.” Unlike P4s, P3s participate in tearing rather than crushing or grinding food. Hlusko and Mahaney (2009) and Hlusko et al. (2011) discovered sub-modularity in baboon maxillary post-canine dentitions, with significant genetic correlations between premolar and molar sizes, and higher correlations within each tooth class. We obtained compatible results based on the shape covariation between human maxillary premolars and molars, reporting a comparable shape covariation pattern, as well as a similar correlation value between upper P4s with both upper M1s (r_1_ = 0.77) and upper M2s (r_1_ = 0.78). Given their distinct functional roles, it is not unexpected to find a lower pairwise correlation between upper P3s and molars. Still, comparable to the P4s, the covariation patterns between upper P3s and both upper molars exhibit a strong morphological resemblance, as the upper molars feature similar morphologies and high correlations with each other (Table 2). Furthermore, Hlusko et al. (2011) reported high genetic correlations between tooth size measurements of upper P3s and P4s for mice and monkeys (length r = 0.84; width r = 0.92), which aligns with our results for the morphological correlation of dentinal crowns in humans. Similarly, for upper molars in a baboon sample, high to moderate values were reported (length r = 0.97; mesial width r = 0.88; distal width r = 76), in agreement with our results for shape correlation in human molars (Hlusko et al., 2011).

In the mandibular postcanine dentition, we observed lower correlation values within the molars than within premolars, and noteworthy, lower than between P4s and M1s. In general, this high pairwise correlation between P4s and M1s in both upper and lower dental arches would support the field theory expressed by Butler (1939). He claimed that all teeth within the postcanine dentition derive from one type of tooth germs under the influence of incomplete pleiotropy and different effects of morphogens on various tooth types. However, it is important to keep in mind that this model stands on primarily phenotypic data, with inconclusive results in subsequent studies aimed at testing Butleŕs hypothesis (Henderson & Greene, 1975; Lombardi, 1975). On the other hand, since P4s and M1s are both engaged in the same function during mastication (Kraus et al., 1969), functional modularity might exert an influence on the morphological covariation of these two dental types as well.

In her study on first molars, Halász (2019) observed a high morphological correlation between the occlusal surfaces of the upper and lower M1s only (r_1_ = 0.85), but a much lower correlation when considering the whole dentinal crown (r_1_ = 0.61) which was analyzed in the current study. Since the pairwise correlation between the upper and lower first molars has been the highest in our study (r_1_ = 0.90), we cannot replicate her results, possibly due to the impact of the sample size on the analyses (n = 18 in Halász’s study; n = 31 in the current study). However, the high correlation of occlusal surfaces of antagonistic teeth is notable, given that these tissues develop following distinct odontogenic pathways. This suggests that apart from developmental integration and pleiotropy, functional integration might underlie the selection of certain trait combinations (correlational selection; Mitteroecker et al., 2012). In contrast to other connective tissue surfaces of the body such as osseous articulations, the only way for antagonistic teeth to alter their shape after initial formation is through wear; this, however, did not affect our analyses of the EDJ (see Introduction and Methods).

Furthermore, our 3D morphological data unraveled a pattern in pairwise correlation values within the postcanine dentition: all of the possible tooth type pairs which included upper M1s showed a high or moderate pairwise correlation value, never lower than r_1_ = 0.67 (non-occluding pair of lower P4 and upper M1), distinguishing upper M1s from the rest of the postcanine dentition. Unlike the other tooth types analyzed in this study, upper M1s do not only correlate strongly in morphology with other teeth in the same dental arch but show high values of morphological covariation with their antagonists and non-occluding tooth types as well. Morita et al. (2016) found M1s exhibiting the least variation in size among upper molars. They considered them the most stable tooth type in morphological terms and suggested upper M1s exhibiting inherent odontogenetic potential. Even though our study is not investigating the inhibitory cascade model (Kavanagh et al., 2007; Morita et al., 2016), our results on the 3D morphology of the upper M1s support the particular status of the upper M1s. Moreover, in the field of dental medicine, first molars are considered pivotal in maintaining the stability of the dental arches throughout growth and development. They play a crucial role in distributing masticatory forces and loads, ensuring the proper function of the stomatognathic system (Slavicek, 2002). Further research is essential to determine whether this is an effect of functional modularity in human evolution, particularly with regard to the upper M1 serving as a key structural pillar for the whole postcanine dentition.

In this study, both the occlusal aspects and the cervical margin of the 3D dentinal crowns were demonstrated by landmark data, capturing the height of the crown as one of the variables in our analysis. Thus, not only the morphology of the occlusal surfaces played a distinctive role in our analyses, but also the general shape-type (see Fig. 3; defined according to the proportions of the crown as tall and narrow or low and broad (Pfneiszl, 2024)) of the dentinal crown was considered. According to Pfneiszl (2024), 87% of individuals from a geographically diverse sample expressed one shape-type consistently across their entire postcanine dentition, thus exhibiting either tall and narrow or low and broad teeth. We detected high pairwise correlations in some non-neighboring and non-occluding tooth type pairs (Supplementary Table S1), highlighting the shared influence of the overall shape-type of the crown. Most of these analyzed tooth type pairs included either the lower premolars or lower M2s, which are known for their morphological variability. Our analysis points to the fact that lower premolars and M2s often exhibited a complex morphological covariation with other teeth, however, this signal was largely masked by the dominant effect of overall crown shape-type. The most extreme value was found between the lower P3s with the upper M2s (r_1_ = 0.88), which covaried almost only with regard to the overall shape-type, while expressing no greater shape changes of the occlusal aspects along the Singular Warp Score 1. On the other hand, distinct tooth types that showed more complex morphological covariation of the dentinal crowns, and thus were not mainly affected by the signal of their shape-type alone, showed low pairwise correlation values (lower P4 and upper M2; r_1_ = 0.44). Methodological limitations, including spatial arrangement of the measurements or geometric dependencies between size-corrected measurements, like Procrustes shape coordinates, may have contributed to these findings by influencing correlation patterns (Mitteroecker, 2009). In related as well as unrelated tooth pairs, particularly if they include an extremely variable tooth type, the general dominance of the overall crown shape-type (tall/narrow vs. low/broad) may lead to a suppression of the signals for the actual shape variation of the occlusal aspect. Confounding factors, possibly affecting our analyses, such as the mode of capturing the geometry of the tooth crowns, need to be considered to fully understand these results. However, since this study is to our knowledge the first one considering morphological covariation in the human postcanine dentition in 3D, obstacles in methodology could be expected and acquire our further attention and tuning.

Nevertheless, given how important teeth are in many fields of natural sciences and medicine, we have relatively little knowledge of odontogenesis. Aspects of cusp pattern development in the interplay of genetic and epigenetic factors still need to be unravelled. Knowledge about the morphological covariation between and within various tooth classes might help supporting future genetic studies. Through our quantitative and qualitative description of the dentinal crown shape covariation of postcanine teeth from various modern human populations, we laid the morphological groundwork for further research on genotype-phenotype associations that is essential to understand odontogenetic processes in mammals, including the genus *Homo*.

## Conclusions

This study extends our understanding of the morphological covariation of human postcanine dentition by analyzing 3D dentinal crown shapes across a geographically diverse sample of modern humans. The results show that upper first molars are a key player in the dental covariation patterns, while developmental integration within dental arches is also evident. Notably, compatibility between antagonistic teeth does not appear to be the primary factor influencing morphological relationships within the postcanine dentition. Furthermore, our results suggest that factors like a shared crown shape-type may dominate the observed covariation patterns, particularly in morphologically variable tooth types like lower premolars. These findings highlight the need for further study of odontogenetic processes and the interplay of genetic and epigenetic factors in tooth development and evolution.

## Acknowledgements

We would like to express our gratitude to Sabine Eggers, Karin Wiltschke-Schrotta, Eduard Winter, and Maria Teschler-Nicola (Natural History Museum, Vienna, Austria), as well as Israel Hershkovitz (Department of Anatomy and Anthropology, Sackler Faculty of Medicine, Tel Aviv University, Israel), for granting us access to the materials. We also thank Martin Dockner (Vienna μCT Lab) for his technical support.

