## Supplementary Information for "Connecting crowns: Analyzing morphological covariation in the modern human postcanine dentition"

^1^Department of Evolutionary Anthropology, University of Vienna, Austria, ^2^Human Evolution and Archaeological Sciences HEAS, University of Vienna, Austria, ^3^Institute of Evolutionary Medicine, University of Zurich, Switzerland,  ^4^Fraunhofer Austria Research GmbH, Klagenfurt, Austria, ^5^Department of Research in Occlusion Medicine, Vienna School of Interdisciplinary Dentistry – VieSID, Klosterneuburg, Austria, ^6^Center for Clinical Research, University Clinic of Dentistry Vienna, Medical University of Vienna, Austria, ^7^Medical Technology Cluster, Business Upper Austria – OÖ Wirtschaftsagentur GmbH, Linz, Austria, ^8^LearnChamp Consulting GmbH & Co KG, Vienna, Austria, ^9^Sandoz GmbH, Kundl, Austria, ^10^Core Facility for Micro-Computed Tomography, University of Vienna, Austria.

**Detailed results of the PLS analyses**

As described in “Material and Methods” of the main text, we performed 2-block Partial Least Squares (2B-PLS) analyses (PLS Mode A; relation between blocks is symmetric) to determine the covariation between tooth types of the human postcanine dentition.

Apart from the morphological changes on the enamel-dentine junction, we observed a general pattern of shape change in dentinal crowns of every tooth type. The tooth types covaried between tall and narrow crowned, and low and broad crowned. Despite a general overlap of the variation between populations, we detected a partial or complete separation of the South American sample from other populations in some of the analyses (Supplementary Fig. S2, S10, S12). However, due to the rather small sample size used in these analyses, and recent literature that points towards the fact that the geographical origin of a modern human individual cannot be detected solely based on the 3D morphological analysis of their dentinal crown (Halász, 2019; Krenn et al., 2019; Šimková et al., 2021, 2024), we consider these findings morphological tendencies.

**A. Morphological covariation of the maxillary teeth**

**Upper P3 – P4**

Both tooth types varied between short-crowned with a mesio-distally broad base and tall-crowned with a mesio-distally narrow base. The occlusal aspect of the short, broad crown type was mesio-distally expanded and square-shaped with a broad central groove. Conversely, tall and slender-crowned teeth had a mesio-distally narrow occlusal aspect with reduced lingual side. Additionally, short-crowned P4s exhibited higher dentinal horn tips compared to tall-crowned premolars.

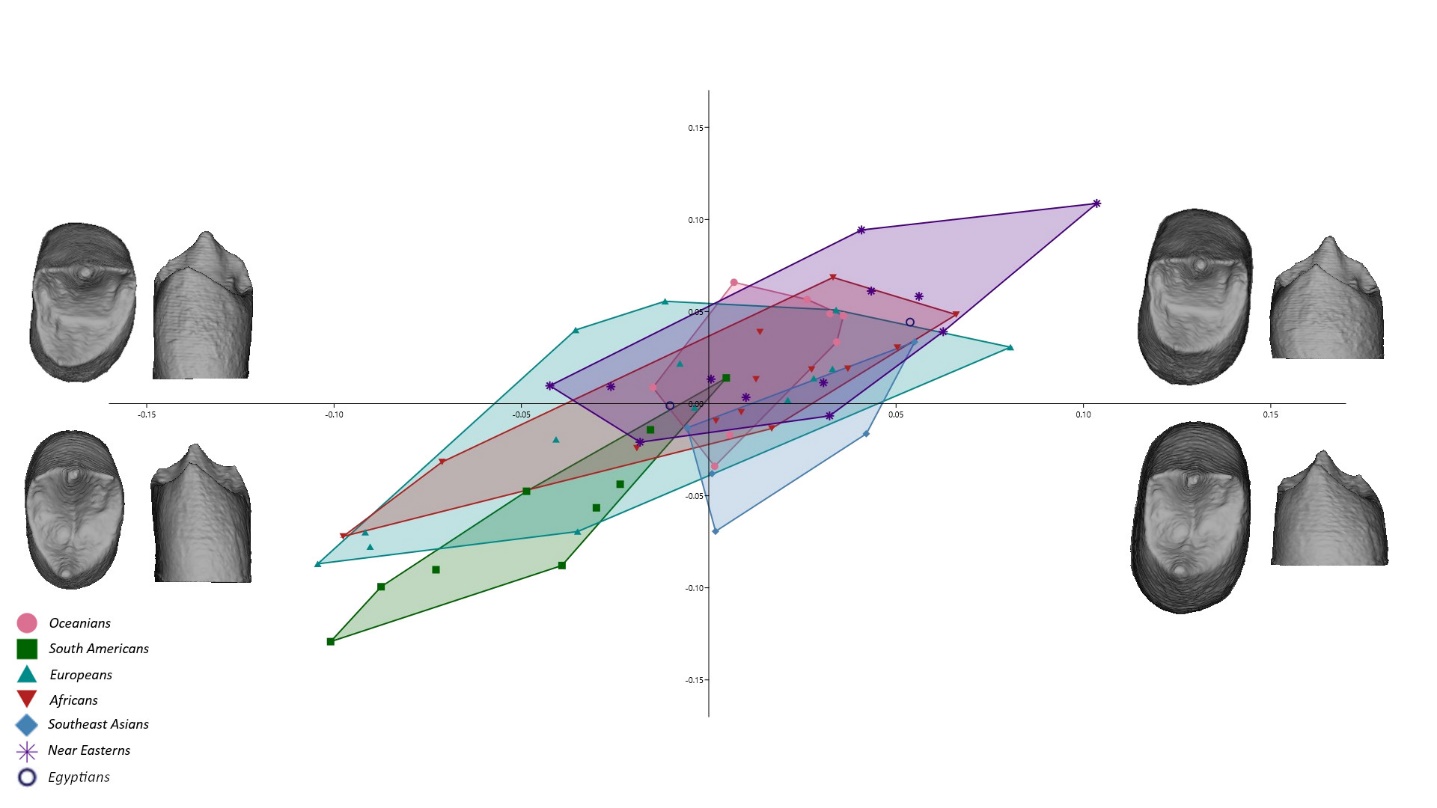

**Supplementary Fig. S1** 2B-PLS plot capturing the covariation of upper third and fourth premolars’ dentinal crowns; the warping shows the real shape variation in occlusal and lingual view at the extremities of the range of distribution.

**Upper P4 – M1**

Main shape changes of both teeth occurred in the morphology of the occlusal aspect, as well as in the change of its size relatively to the base of the tooth. Both dental types varied between large-based with a constricted occlusal aspect and small-based with expanded occlusal aspect. In the latter, the premolar’s buccal part of the occlusal aspect expanded mesio-distally and buccally, with higher buccal cusp relatively to the lingual cusp. The occlusal lingual side was mesio-distally reduced relatively to the buccal part. The molars showed a variable relationship between the trigon and the talon, with a relative expansion and reduction of the hypocone.

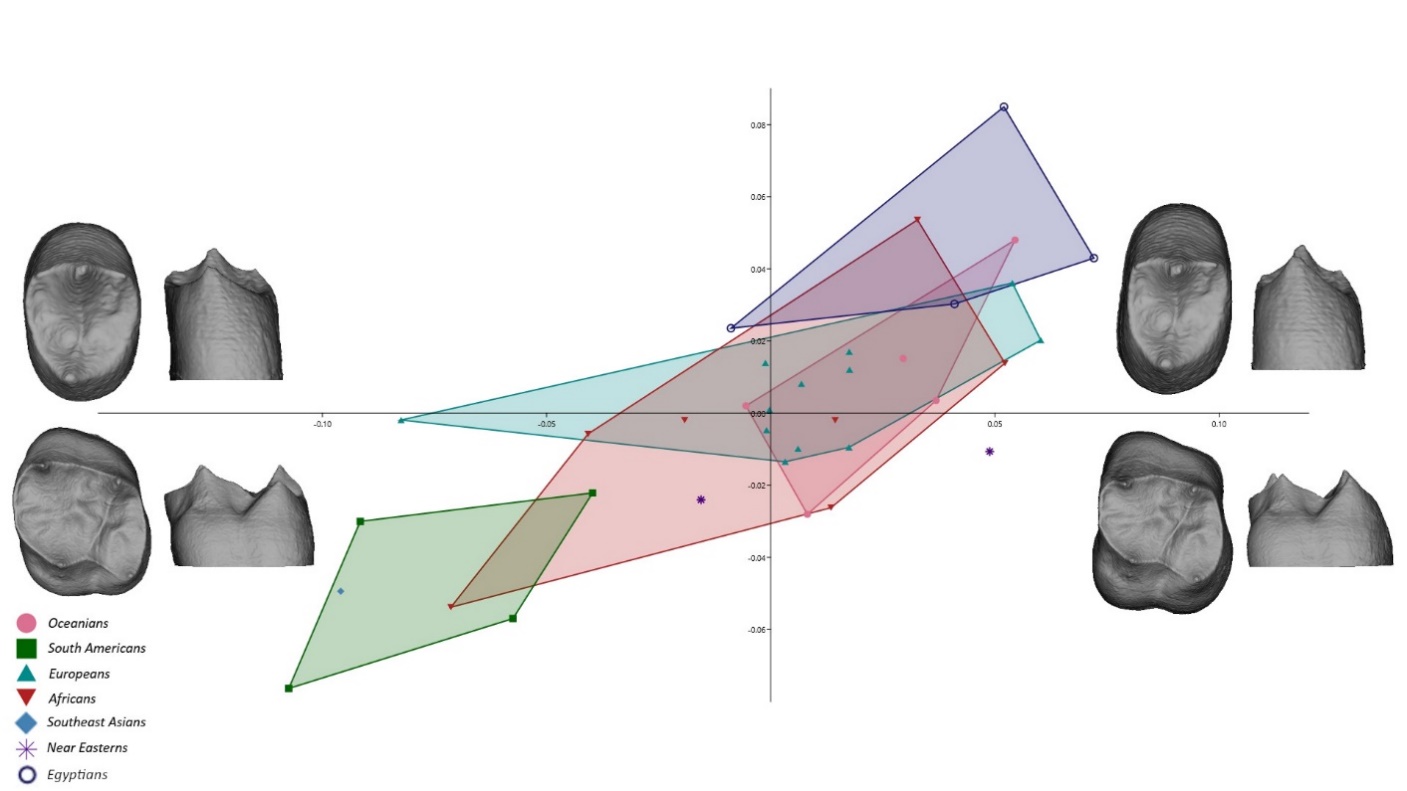

**Supplementary Fig. S2** 2B-PLS plot capturing the covariation of upper fourth premolars’ and first molars’ dentinal crowns; the warping shows the real shape variation in occlusal and lingual view at the extremities of the range of distribution.

**Upper M1 – M2**

The upper M1s and M2s covaried in the same manner. Their morphology varied across the sample between low-crowned with expanded base and relatively large hypocone in comparison to the trigon, and tall-crowned with reduced base, with bucco-lingually expanded trigon and relatively reduced hypocone.

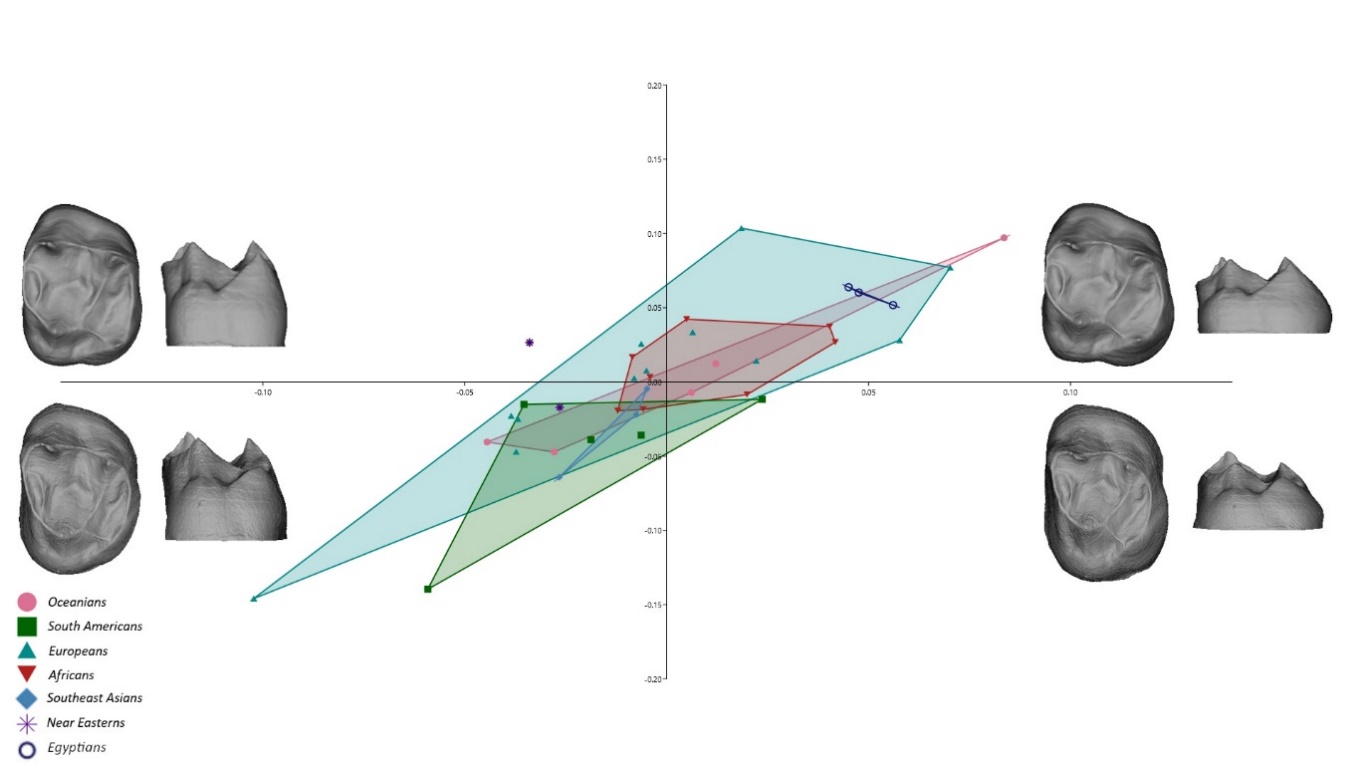

**Supplementary Fig. S3** 2B-PLS plot capturing the covariation of upper first and second molars’ dentinal crowns; the warping shows the real shape variation in occlusal and lingual view at the extremities of the range of distribution.

**B. Morphological covariation of the mandibular teeth**

**Lower P3- P4**

Similar to upper jaw, the covariation analysis between lower premolars reflected shape change in both tooth types mainly between short, broad- crowned teeth with expanded, squared occlusal aspect, resulting from disto-lingual shift of the lingual cusp, and tall, narrow- crowned teeth with mesially shifted lingual cusp and relatively smaller mesial fovea of the central fossa in comparison to the distal fovea.

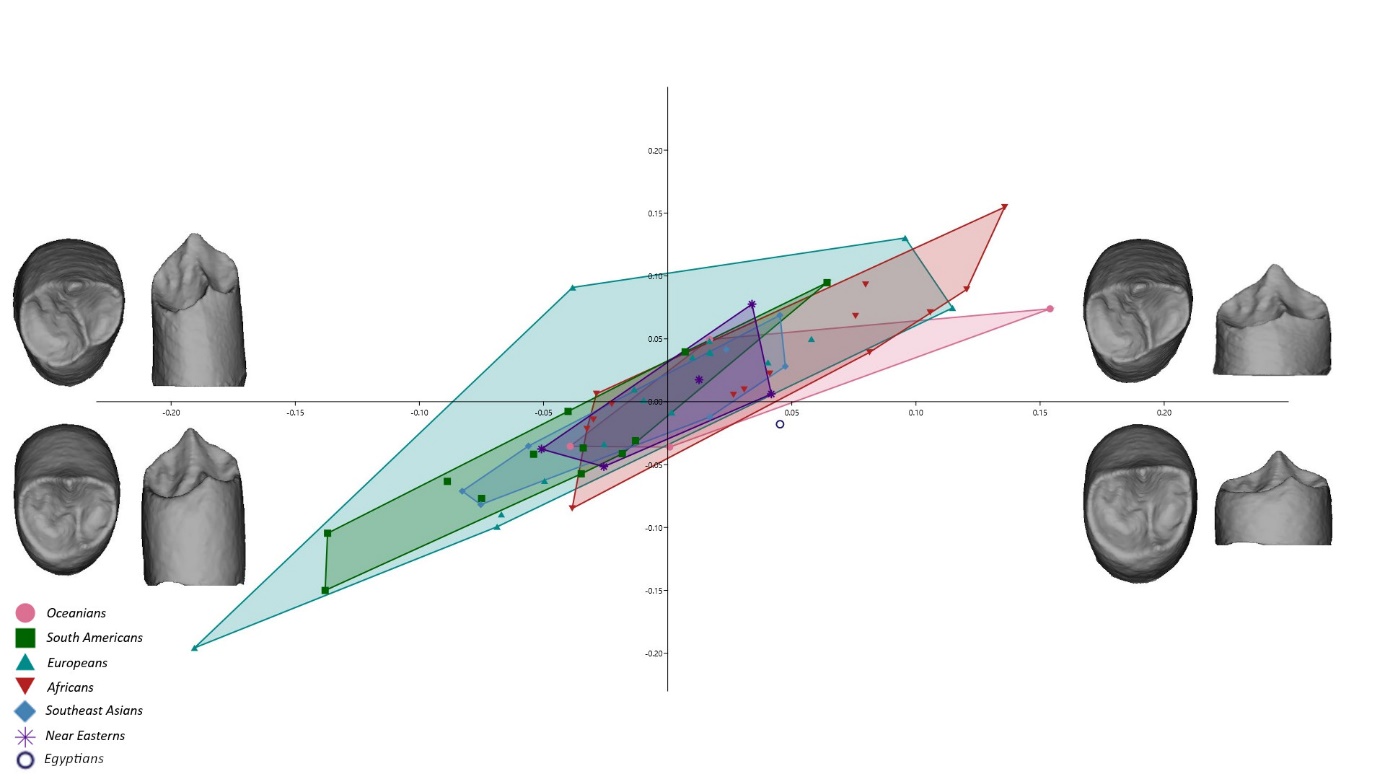
**Supplementary Fig. S4** 2B-PLS plot capturing the covariation of lower third and fourth premolars’ dentinal crowns; the warping shows the real shape variation in occlusal and lingual view at the extremities of the range of distribution.

**Lower P4 – M1**

Covariation analysis of lower P4s’ and M1s’ dentinal crowns, reflected main shape change between tall and narrow crowns, and short and broad crowns in both dental types. The expansion and reduction of the base was relatively consistent with the expansion and reduction of the occlusal aspect. Tall- crowned P4s featured relatively bigger increase in height of the buccal cusp in comparison to the short- crowned teeth.

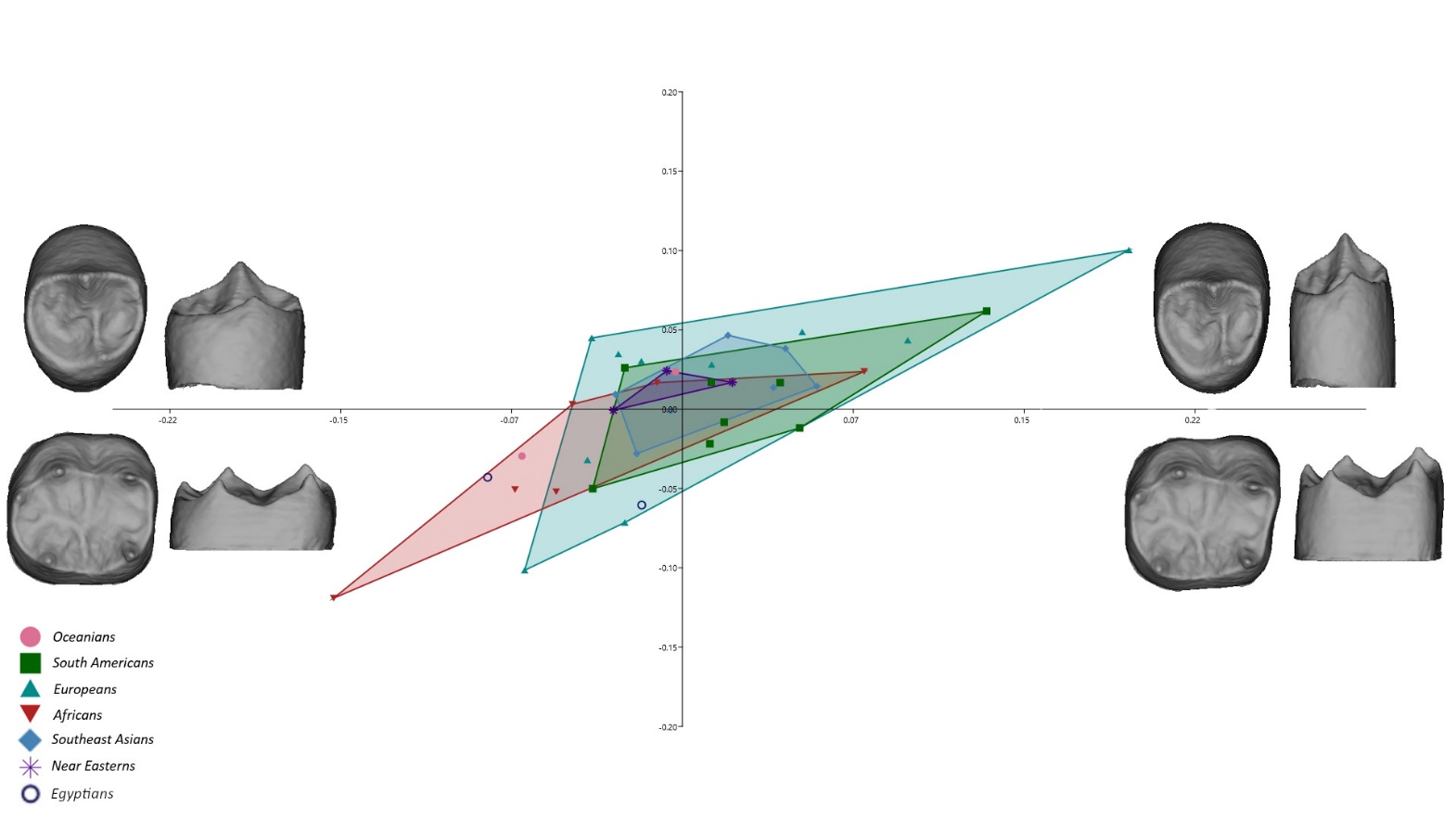

**Supplementary Fig. S5** 2B-PLS plot capturing the covariation of lower fourth premolars’ and first molars’ dentinal crowns; the warping shows the real shape variation in occlusal and lingual view at the extremities of the range of distribution.

**Lower M1 – M2**

The crowns of both tooth types varied mainly between short and broad crowns, with short cusps, and a barrel shaped base and occlusal aspect, and tall crowns with tall cusps, bucco-lingually constricted narrow base and occlusal aspect, and disto-lingually reduced occlusal aspect in M2s.

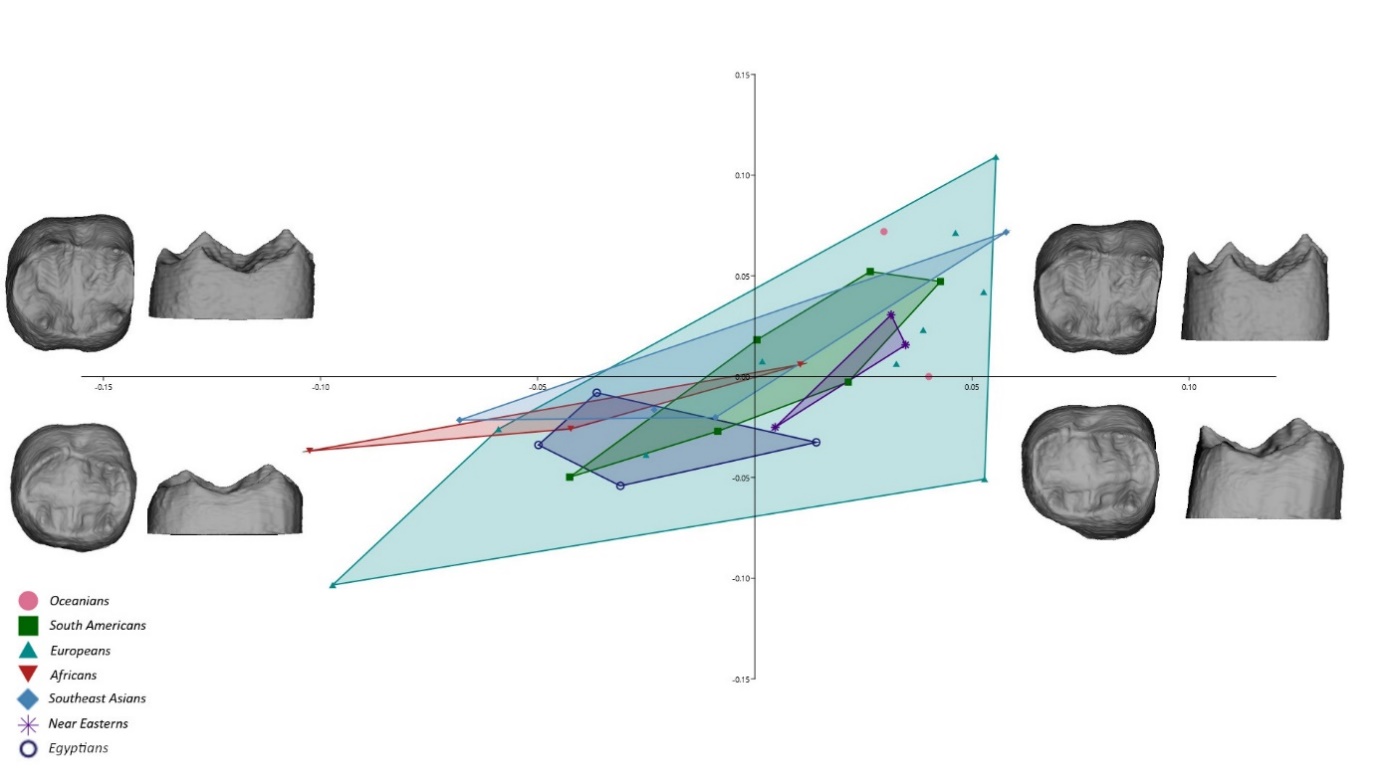
**Supplementary Fig. S6** 2B-PLS plot capturing the covariation of lower first and second molars’ dentinal crowns; the warping shows the real shape variation in occlusal and lingual view at the extremities of the range of distribution.

**C. Morphological covariation between antagonists**

**Lower P3 – Upper P3**

The crowns of antagonistic P3s covaried mainly between tall and narrow-crowned, and short and broad-crowned. The first type showed reduction of the distal and disto-lingual aspect in the upper P3, as well as in lower P3s that additionally showed a relatively taller buccal cusp and lingual shifting of the lingual cusp.

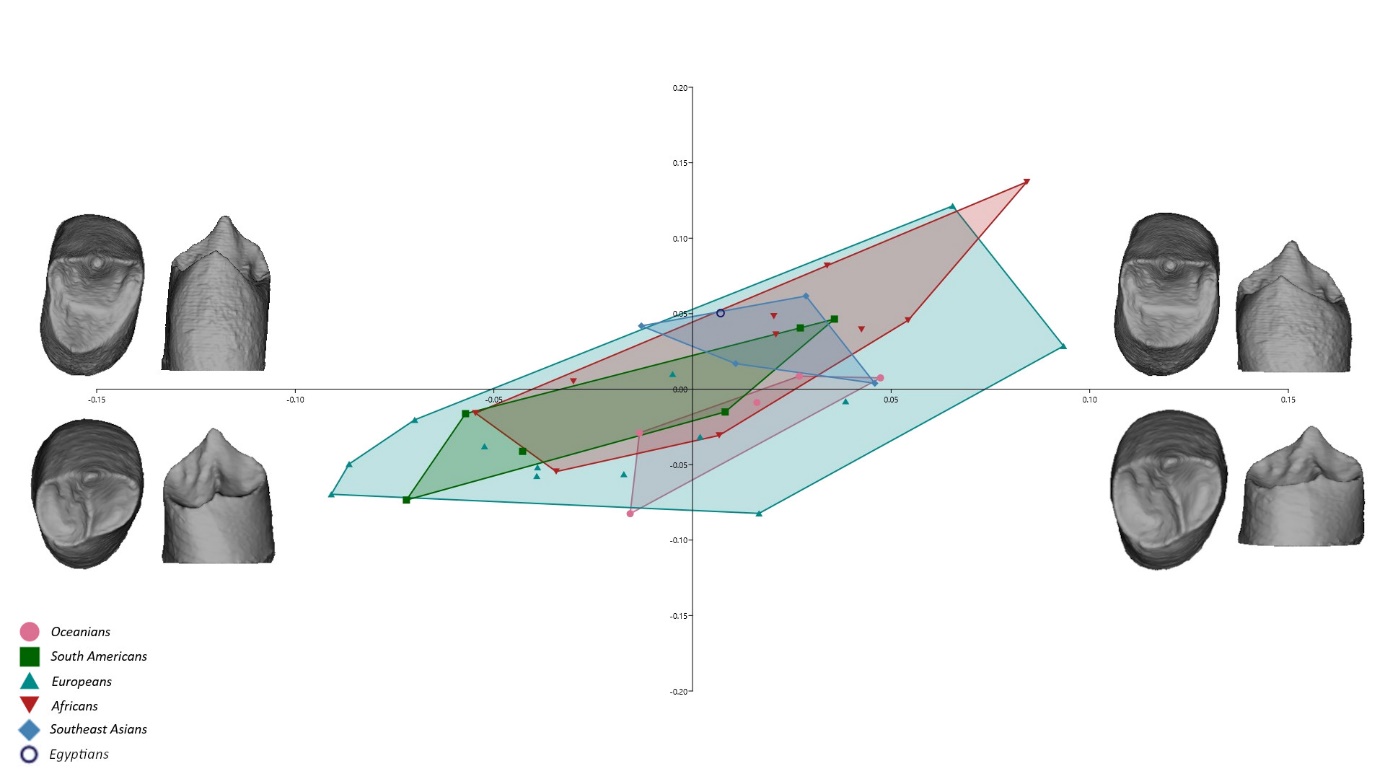

**Supplementary Fig. S7** 2B-PLS plot capturing the covariation of upper and lower third premolars’ dentinal crowns; the warping shows the real shape variation in occlusal and lingual view at the extremities of the range of distribution.

**Lower P4 – Upper P4**

Similar to P3s, in P4s shape changes occurred mainly between tall and narrow-crowned and short and broad-crowned teeth. The buccal cusp of the lower P4s was variably expanded with respect to the rest of the crown.

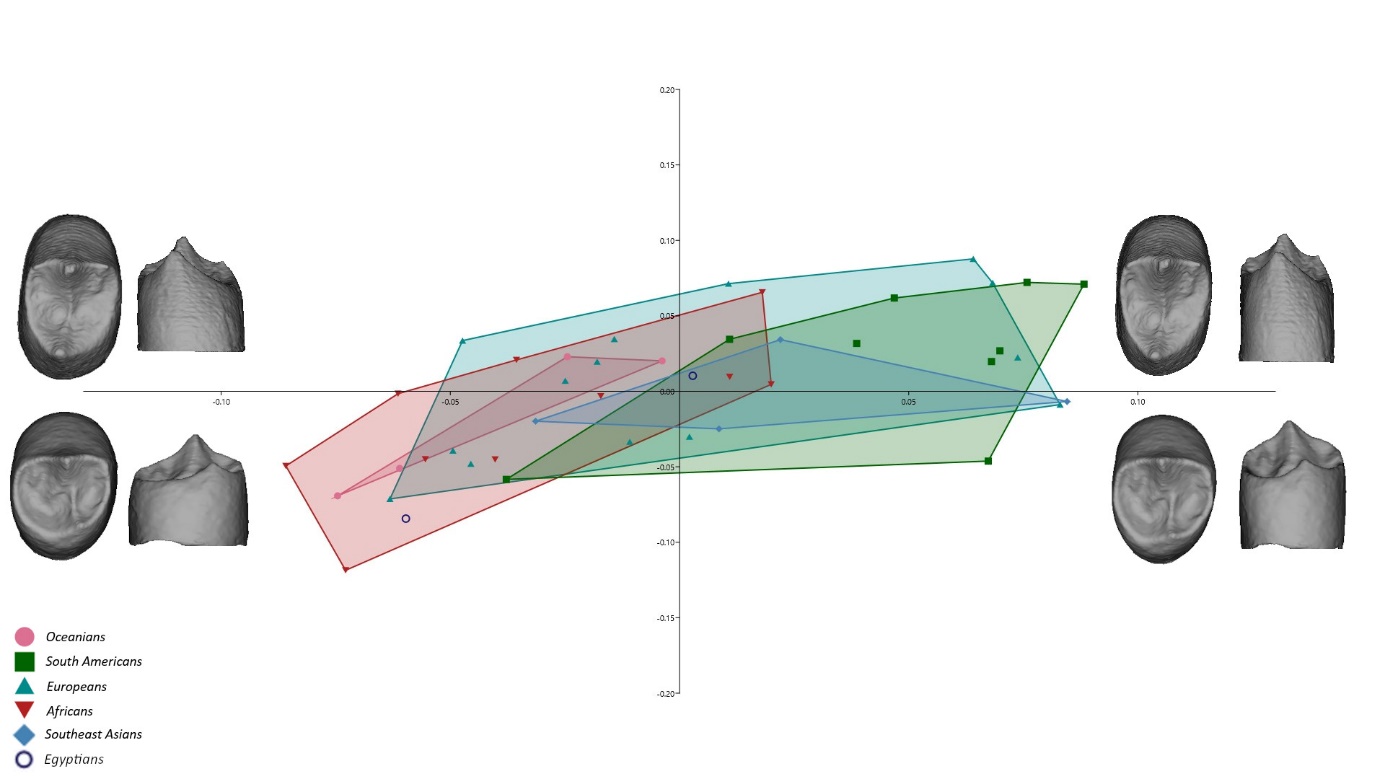

**Supplementary Fig. S8** 2B-PLS plot capturing the covariation of upper and lower fourth premolars’ dentinal crowns; the warping shows the real shape variation in occlusal and lingual view at the extremities of the range of distribution.

**Upper P3 – Lower P4**

The premolars covaried mainly between tall and narrow-crowned to short and broad-crowned, the latter presenting with expanded distal side of the occlusal aspect creating a squared shape in both P3s and P4s. Tall and narrow crowned lower P4s featured a taller buccal cusp and disto-lingually shifted lingual cusp.

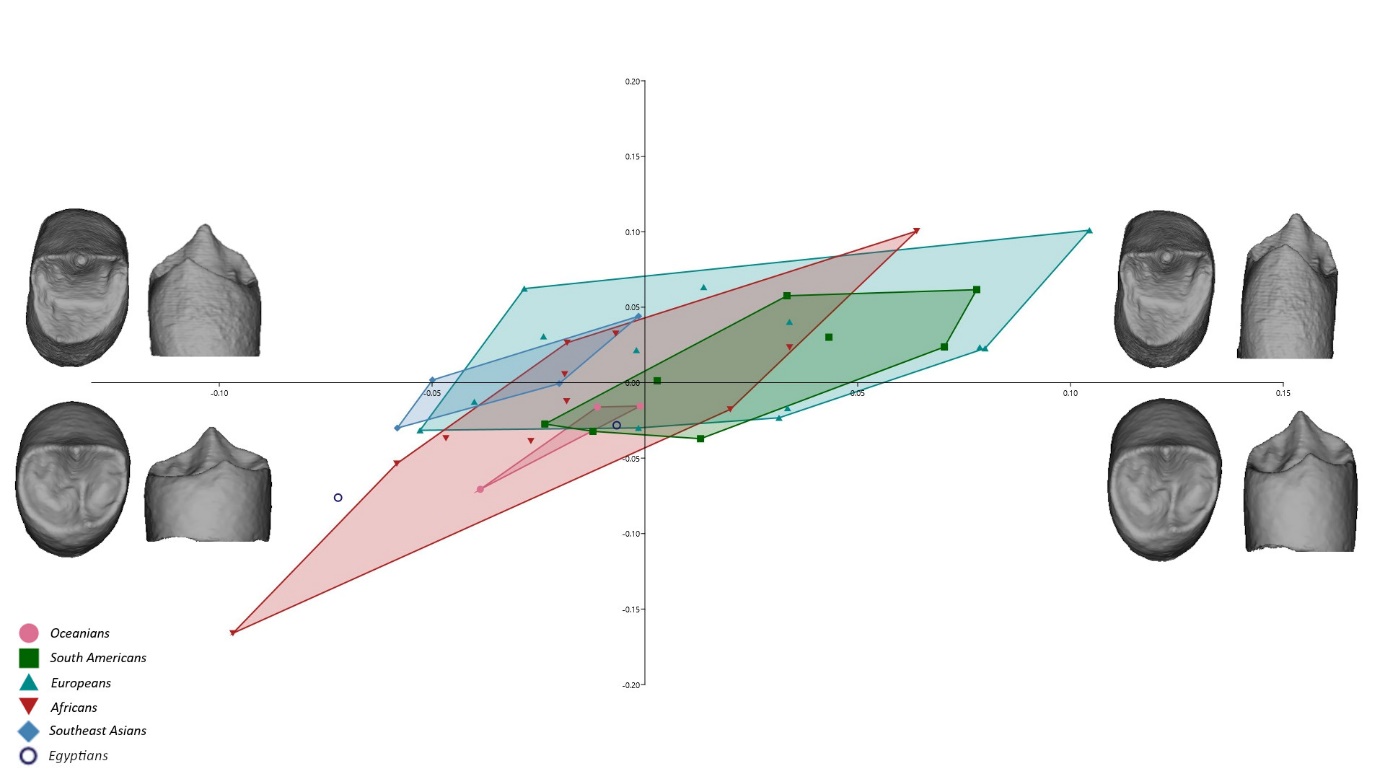

**Supplementary Fig. S9** 2B-PLS plot capturing the covariation of upper third and lower fourth premolars’ dentinal crowns; the warping shows the real shape variation in occlusal and lingual view at the extremities of the range of distribution.

**Upper P4 – Lower M1**

The morphological covariation occurred mainly in the height of the dental crowns and in their basal relative size. Shape variation of the occlusal aspect was mild and driven by the mesio-distal shifting of the lingual cusp in P4s, and relative inclination of the molar horn tips either towards or away from each other.

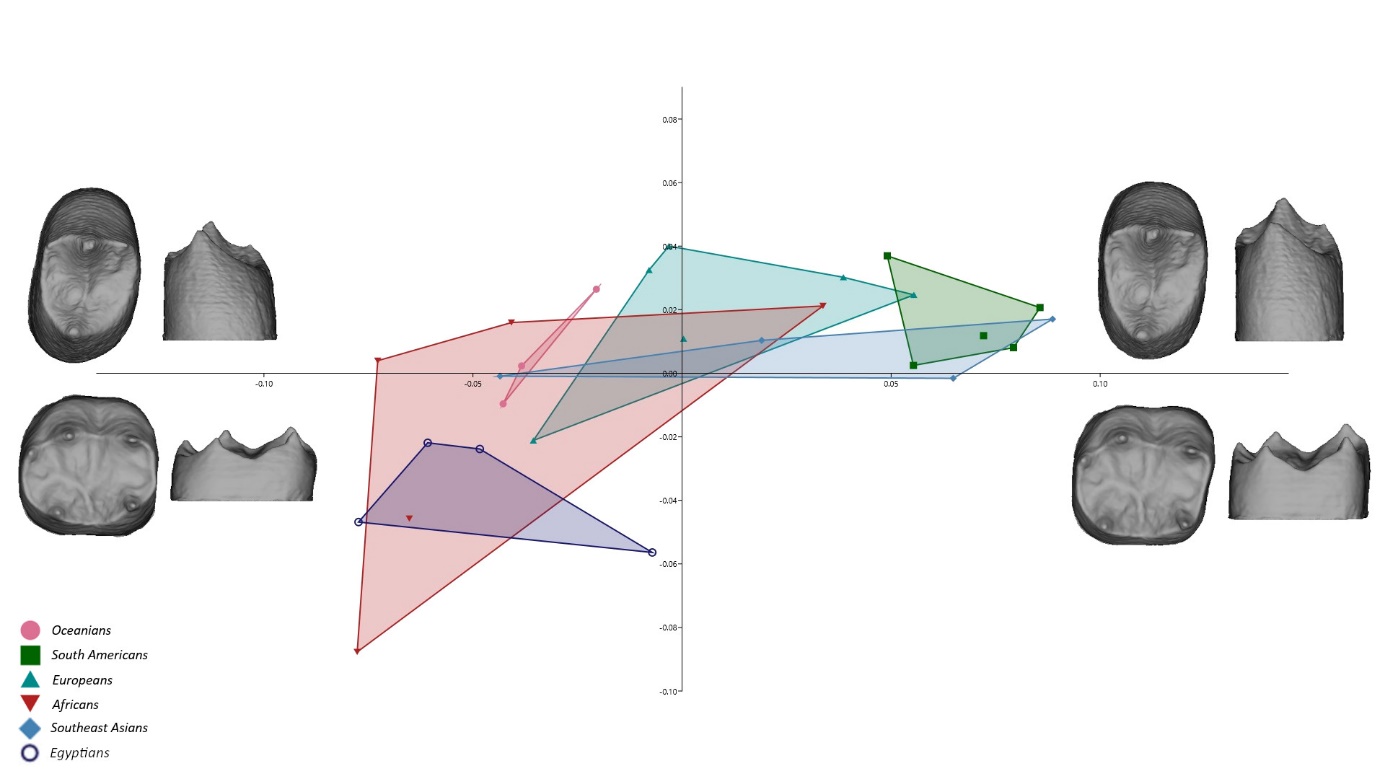

**Supplementary Fig. S10** 2B-PLS plot capturing the covariation of upper fourth premolars’ and lower first molars’ dentinal crowns; the warping shows the real shape variation in occlusal and lingual view at the extremities of the range of distribution.

**Upper M1 – Lower M2**

The shape changes were evident in the marked relative expansion and reduction of the distal and disto-lingual aspect of the dentinal crown reflected in the general presentation of the crown as broader or narrower.

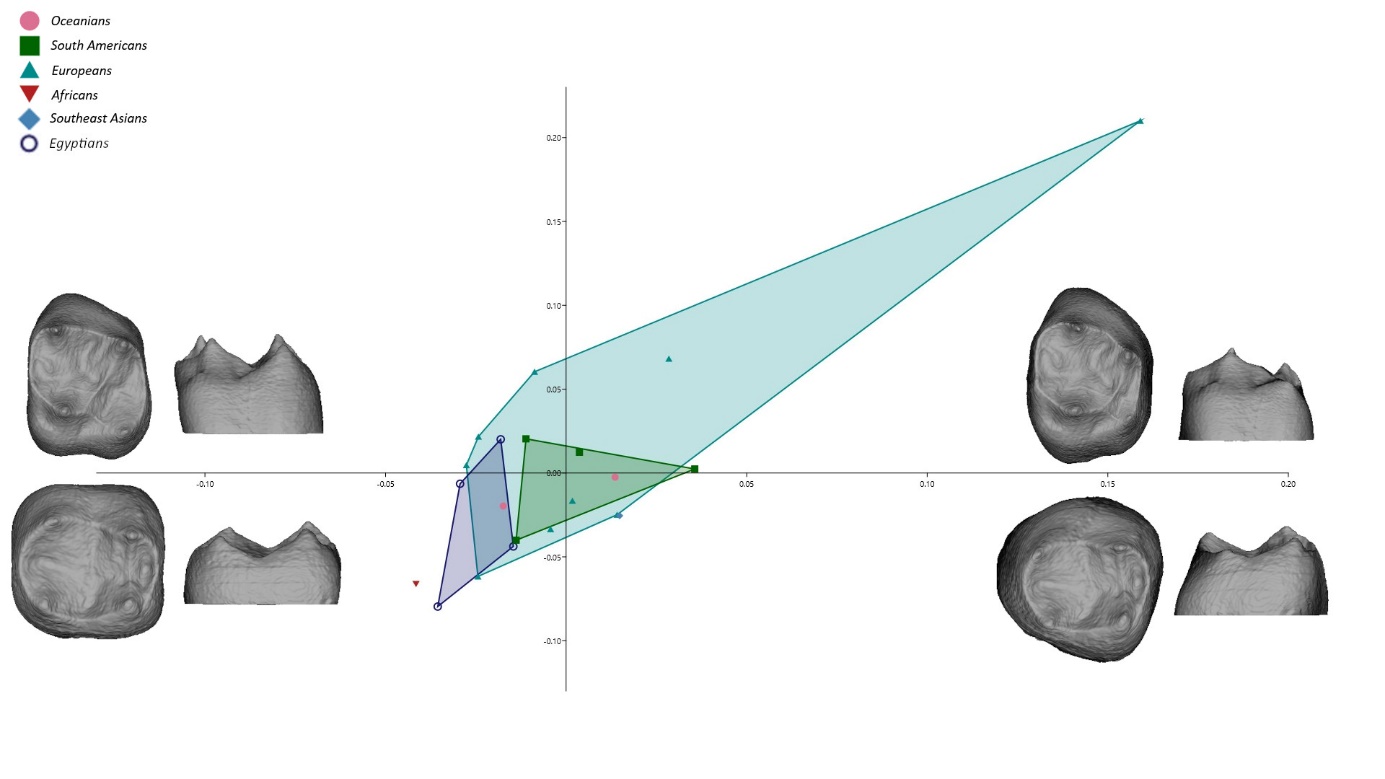

**Supplementary Fig. S11** 2B-PLS plot capturing the covariation of upper first and lower second molars’ dentinal crowns; the warping shows the real shape variation in occlusal and lingual view at the extremities of the range of distribution.

**Lower M2 – Upper M2**

The antagonistic M2s covaried mainly in the height of the crown and shape of the crown base. Tall crowns were mesio-distally constricted in upper M2s, featuring a relatively reduced base in comparison to the occlusal aspect that was more squared, while tall lower molar showed a bucco-lingually constricted base and expanded distal part of the occlusal aspect. Short crowns presented a large base relative to the occlusal aspect, mesio-distally expanded in upper molars, with lingually shifted hypocone. Lower molars showed a bucco-lingually, as well as mesio-lingually expanded base, featuring a reduction of the distal part of the occlusal aspect.

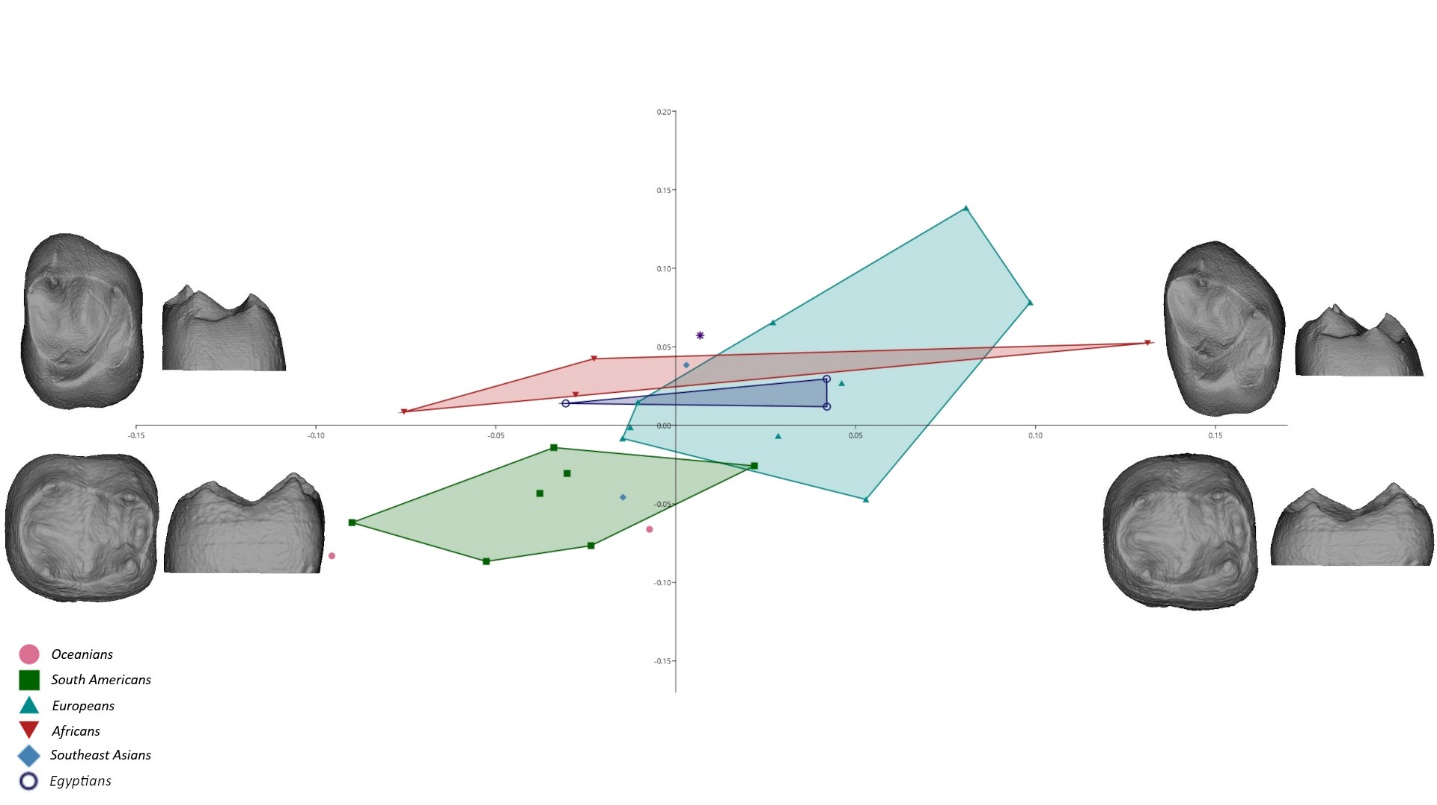
**Supplementary Fig. S12** 2B-PLS plot capturing the covariation of upper and lower second molars’ dentinal crowns; the warping shows the real shape variation in occlusal and lingual view at the extremities of the range of distribution.

**D. Morphological covariation between non-occluding tooth types**

**Lower P3 – Upper P4**

The results of the 2B-PLS analysis showed morphological covariation between lower P3s and upper P4s to be mainly between tall and narrow-crowned type and short and broad type of the crowns. In the upper premolars, the occlusal aspect varied from squared in short crowns, to narrow and pointy, due to the lingual shifting of the lingual cusp in taller crowns. In the lower premolars, the position of the lingual cusp varied between mesially and lingually shifted, resulting in varying expansion of the mesial fossa.

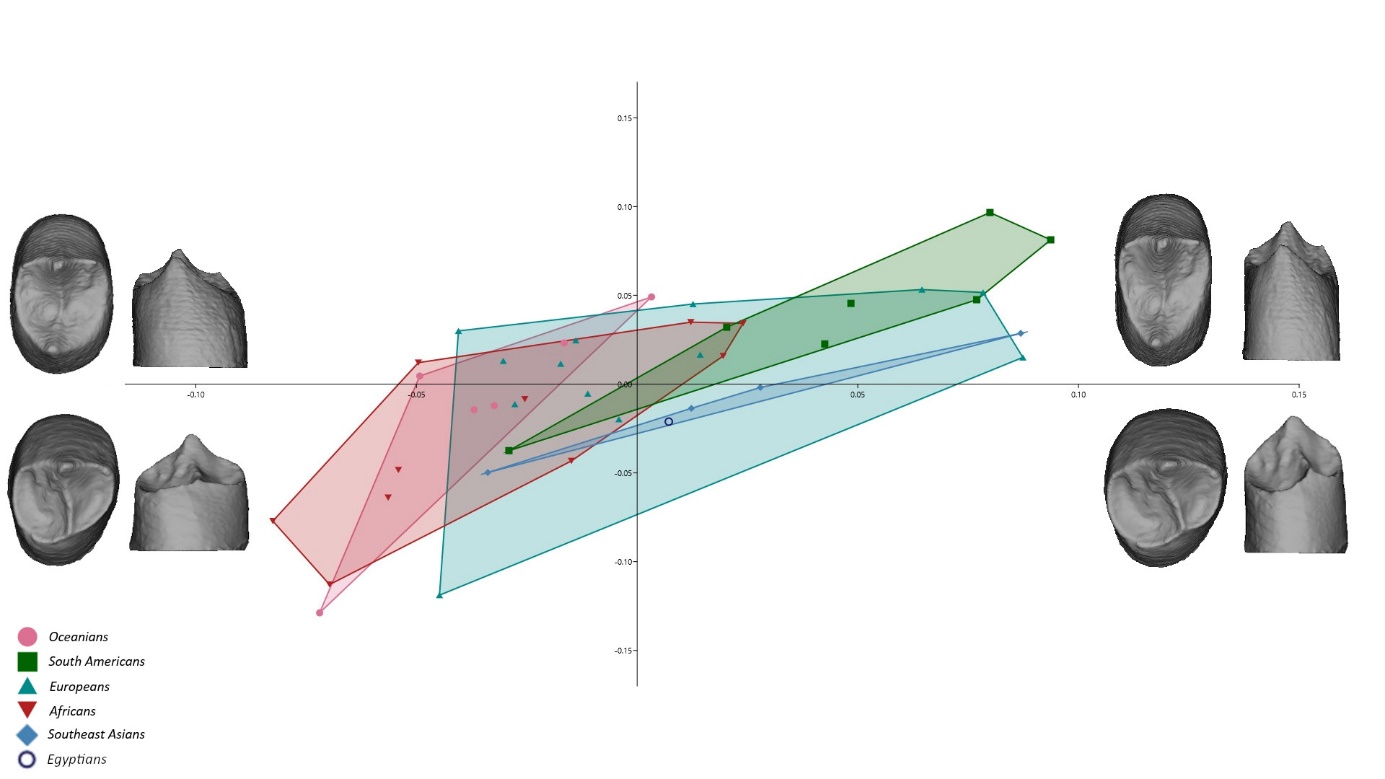
**Supplementary Fig. S13** 2B-PLS plot capturing the covariation of lower third and upper fourth premolars’ dentinal crowns; the warping shows the real shape variation in occlusal and lingual view at the extremities of the range of distribution.

**Lower P4 – Upper M1**

Lower P4s’ and upper M1s’ crowns covaried between tall and narrow with relatively reduced mesio-lingual side of the occlusal aspect in premolars, and disto-lingual side in molars, and short and broad crowns with relative expansion of the same aspects.

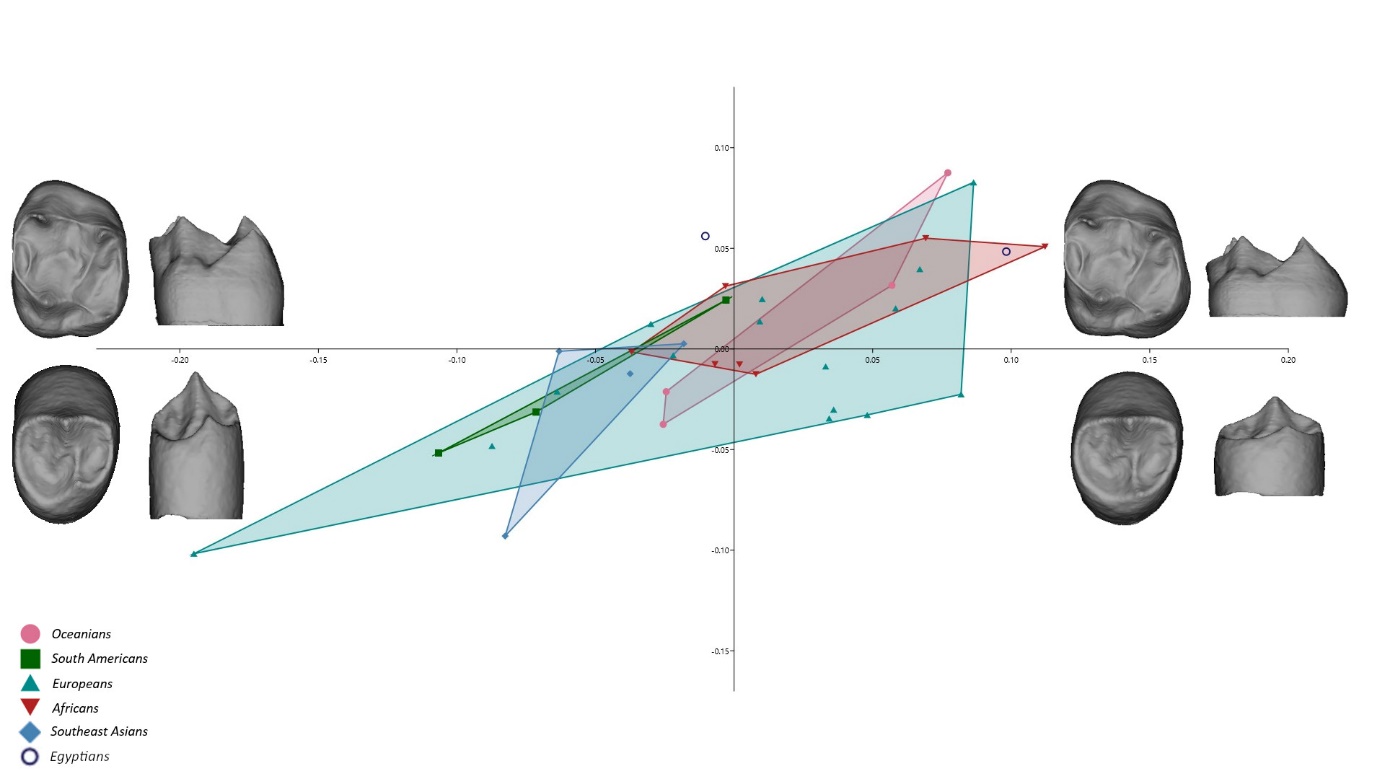

**Supplementary Fig. S14** 2B-PLS plot capturing the covariation of lower fourth premolars’ and upper first molars’ dentinal crowns; the warping shows the real shape variation in occlusal and lingual view at the extremities of the range of distribution.

**Lower M1 – Upper M2**

Both tooth types covaried between tall crowns with narrow occlusal aspects and lingually positioned hypocone/hypoconulid, and short crowns with broad base and buccally positioned hypocone/hypoconulid.

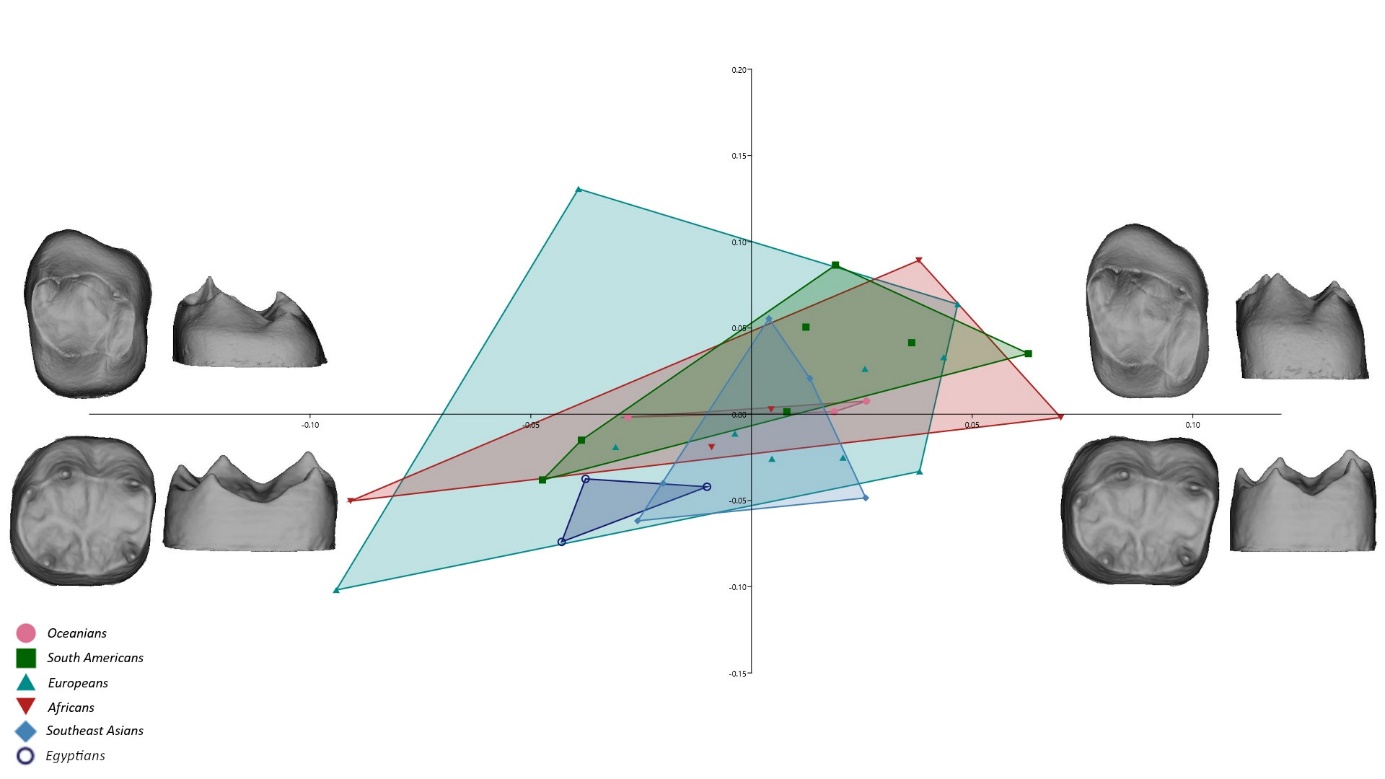

**Supplementary Fig. S15** 2B-PLS plot capturing the covariation of lower first and upper second molars’ dentinal crowns; the warping shows the real shape variation in occlusal and lingual view at the extremities of the range of distribution.

**E. Pairwise correlation values indicated by occluding/neighboring and non-occluding/non-neighboring status**

| **Supplementary Table S1. Colormap of the pairwise correlation values** resulting from 2-Block Partial Least Squares Analyses. (L = lower; U = upper; P3 = third premolars: P4 = fourth premolars; M1 = first molars; M2 = second molars).  Blue colors indicate occluding or neighboring tooth type pairs; orange colors indicate non-occluding and non-neighboring tooth-type pairs. In both instances, darker colors indicate stronger correlations. | | | | | | | | |
| --- | --- | --- | --- | --- | --- | --- | --- | --- |
|  | LP3 | LP4 | LM1 | LM2 | UP3 | UP4 | UM1 | UM2 |
| LP3 | 1 | 0.89 | 0.76 | 0.74 | 0.71 | 0.75 | 0.77 | 0.88 |
| LP4 |  | 1 | 0.79 | 0.75 | 0.70 | 0.65 | 0.73 | 0.44 |
| LM1 |  |  | 1 | 0.73 | 0.61 | 0.58 | 0.90 | 0.50 |
| LM2 |  |  |  | 1 | 0.63 | 0.66 | 0.82 | 0.64 |
| UP3 |  |  |  |  | 1 | 0.81 | 0.67 | 0.70 |
| UP4 |  |  |  |  |  | 1 | 0.77 | 0.78 |
| UM1 |  |  |  |  |  |  | 1 | 0.83 |
| UM2 |  |  |  |  |  |  |  | 1 |
